# The Impact of Model Misspecification on Phylogenetic Network Inference

**DOI:** 10.1101/2022.10.24.513600

**Authors:** Zhen Cao, Meng Li, Huw A Ogilvie, Luay Nakhleh

**Affiliations:** Department of Computer Science, Rice University, Houston, TX, USA; Department of Statistics, Rice University, Houston, TX, USA; Department of Biosciences, Rice University, Houston, TX, USA

## Abstract

The development of statistical methods to infer species phylogenies with reticulations (species networks) has led to many discoveries of gene flow between distinct species. These methods typically assume only incomplete lineage sorting and introgression. Given that phylogenetic networks can be arbitrarily complex, these methods might compensate for model misspecification by increasing the number of dimensions beyond the true value. Herein, we explore the effect of potential model misspecification, including the negligence of gene tree estimation error (GTEE) and assumption of a single substitution rate for all genomic loci, on the accuracy of phylogenetic network inference using both simulated and biological data. In particular, we assess the accuracy of estimated phylogenetic networks as well as test statistics for determining whether a network is the correct evolutionary history, as opposed to the simpler model that is a tree.

We found that while GTEE negatively impacts the performance of test statistics to determine the “tree-ness” of the evolutionary history of a data set, running those tests on triplets of taxa and correcting for multiple testing significantly ameliorates the problem. We also found that accounting for substitution rate heterogeneity improves the reliability of full Bayesian inference methods of phylogenetic networks, whereas summary statistic methods are robust to GTEE and rate heterogeneity, though currently require manual inspection to determine the network complexity.

## 2 Introduction

Our understanding of evolutionary history is limited by the models used to represent those histories. The most common model is a phylogenetic *tree*, which may be used to model the histories of genes (by which we mean discrete parts of a genome called loci) and species. When a phylogenetic tree is used to model the history of a set of genes related by common descent, it is known as a gene tree. When used to model species histories and hence known as species trees, branching events represent instances of speciation, or the splitting of one ancestral species into descendant species. Species trees by themselves do not permit reticulation, instead building on an assumption that once a species is divided, no gene flow (where individuals from different contemporaneous species may produce offspring) between the divided populations will occur subsequently. Species tree inference despite the presence of gene flow has been explored (Davidson et al., 2015; Roch & Snir, 2013); in addition to the fact that such a species tree gives an incomplete picture of the evolutionary history, it could also be an incorrect tree (Cao et al., 2019; Solís-Lemus et al., 2016; Zhu et al., 2016). Forms of gene flow that may occur are introgression (the transmission of genes from one species to another) and hybrid speciation (where the offspring of parents from different species form a new species). As a result, the discovery of many instances of reticulate evolution was delayed until the first methods were developed to detect gene flow.

The development of methods to infer reticulate evolution among eukaryotic nuclear genomes is particularly challenging because incomplete lineage sorting (ILS) causes incongruence of gene and species phylogenies even in the complete absence of reticulations (Nakhleh, 2013). Gene duplication and loss alone or in concert with ILS may also result in incongruence (Rasmussen & Kellis, 2012). Nevertheless, methods which discriminate between these processes have been developed—for example, the ABBA-BABA statistical test enabled the detection of introgression from Neanderthals into modern humans (Green et al., 2010).

The identification of introgression in eukaryotes motivated the development of combinatorial and statistical methods that use phylogenetic *networks* to model the evolutionary history of species (henceforth *species networks*; Flouri et al., 2020; Wen & Nakhleh, 2018; Wen, Yu, & Nakhleh, 2016; Yu et al., 2013; Yu et al., 2014; Zhang et al., 2018; Zhu & Nakhleh, 2018; Zhu et al., 2018). Such methods generalize the multispecies coalescent (MSC) model, which specifies a probability distribution over gene trees given a species tree and a demographic model. The generalized model instead uses a species network and is known as the multispecies network coalescent (MSNC) model. MSNC methods enabled the discovery of more instances of introgression (Fontaine et al., 2015; Marcussen et al., 2014), demonstrating the importance of tractable and reliable methods for detecting and characterizing reticulate evolution.

One class of MSC and MSNC methods is multilocus methods, which use alignments of genes as input. These methods assume recombination is frequent between genes and limited within them, so that a single gene tree can be inferred for individual alignments, and the histories of different alignments are independent samples from the underlying distribution of gene trees. These methods may be further divided into summary and full methods. Summary methods use previously inferred gene trees to estimate the species phylogeny, whereas full methods jointly estimate the species phylogeny and gene trees in a single step.

In the analysis of real data, when gene tree discordance is observed, both MSC and MSNC methods can be applied to obtain the species phylogeny (Edelman et al., 2019; Morales-Briones et al., 2021). However MSC methods will converge on a species tree even if a species network is a better representation of reality, and we hypothesized that MSNC methods may due to model misspecification converge on a network even if a species tree is a better representation of reality. Therefore the species phylogeny estimated by MSC methods, and possibly even those estimated by MSNC methods, cannot tell us whether the best-fitting model of evolution for a group of species is a tree or network.

One approach to determine the best-fitting model is the use of previously proposed statistical methods which apply Pearson’s *χ*^2^ test (Degnan & Rosenberg, 2009) to examine the goodness-of-fit of the MSC by comparing the expected and observed frequencies of gene tree topologies (Cai & Ané, 2021; Stenz et al., 2015). As we will demonstrate, these statistical methods can be misled in practice with inevitable gene tree estimation error (GTEE) due to the fact that gene trees are inferred from sequence data.

Although full Bayesian methods are designed with the inherent ability to detect introgression on their own through the incorporation of prior knowledge, they may be faced with model misspecification caused by heterogeneity of substitution rates, which are the rates genes evolve at, and until now the effects of substitution rate heterogeneity on network inference methods have not been explored. Substitution rates are suspected to vary greatly between loci (Duret & Mouchiroud, 2000), changing the branch lengths of gene trees when measured in the expected number of substitutions. Summary methods, such as `InferNetwork_ML` (Yu et al., 2014), are usually set to ignore branch lengths, and so should be robust to differences in true branch lengths, although perhaps not to the increased error which is inversely correlated with branch length.

However, the full method `MCMC_SEQ` (Wen & Nakhleh, 2018) previously assumed all loci evolve at the same substitution rate and may be misled since the species network probability is dependent on coalescent times within each gene tree which are scaled by the substitution rate at each locus. This is a particular concern when reticulations are permitted by the model and method, since spurious reticulations may be added to the species network in order to account for the apparently (but not actually) different coalescent times.

Herein we seek to understand the effect of model misspecification on phylogenetic network inference. We ask two questions regarding the model misspecification from GTEE and rate heterogeneity. First, can test statistics detect introgression correctly when gene trees are inferred? Second, can summary methods and full network inference methods infer the species phylogeny accurately in the presence of substitution rate variation?

To address the first question, we perform a simulation study to assess several statistical tests for their ability to determine the goodness-of-fit of the MSC model to multi-locus alignment data, which gene trees are estimated from with error. We refined the test statistics by developing a “triplet multitest” that applies test statistics to every set of three taxa, then applies a correction for multiple testing.

To address the second question, we conduct a simulation study to understand their behavior and susceptibility to error. We have also enhanced `MCMC_SEQ` by incorporating support for varying substitution rates with a fixed mean and an implied flat Dirichlet prior by adding a delta exchange operator (DEO) to that method. Henceforth, in the context of simulation and inference, we will refer to this as the “Dirichlet rates” (DR) model, as opposed to a single rate (SR) for all loci. Both the DEO and DR model of substitution rate variation were originally introduced in BEAST (Drummond & Rambaut, 2007). We show that allowing for substitution rate variation dramatically improves the accuracy of species network inference when rate variation is present in the data, and does not meaningfully reduce the accuracy of inference even when all loci have the same rate.

By applying both the novel triplet multitest statistical method and the DEO-enhanced version of `MCMC_SEQ`, we show that previously reported reticulations in *Anopheles* mosquitoes are likely to reflect the reality of that clade, while at least some of the previously reported reticulations in *Heliconius* butterflies may be artifacts of model misspecification.

Our implementation of the Dirichlet per-locus rates model is now available in `PhyloNet` (Than et al., 2008; Wen et al., 2018), a software package for phylogenetic inference, open source on GitHub https://github.com/NakhlehLab/PhyloNet. Code implemented for our simulations, analysis and visualization is also available on GitHub: https://github.com/NakhlehLab/Model-Misspecification. The empirical data sets of *Anopheles* and *Heliconius* data we used were retrieved from Dryad: https://datadryad.org/stash/dataset/doi:10.5061/dryad.tn47c and https://datadryad.org/stash/dataset/doi:10.5061/dryad.b7bj832 respectively.

## 3 Methods

### 3.1 Simulated multi-locus data

For model species phylogenies, we used the displayed tree and network of the evolutionary history of anopheline mosquito species (Fontaine et al., 2015) restricted to *An. coluzzii*, *An. gambiae*, *An. quadriannulatus*, *An. arabiensis*, and *An. melas* (Figure 1).

**Figure 1:**
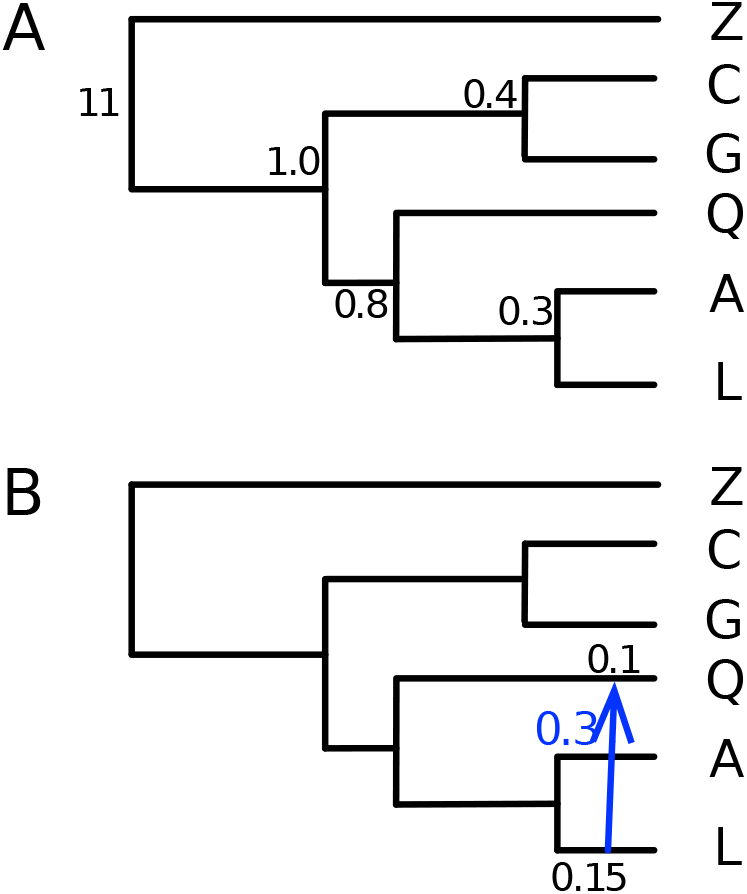
Exemplar phylogenies used in our simulation study. The two species phylogenies were chosen from a displayed subtree of anopheline mosquitoes (A) and a displayed subnetwork of the same taxa (B). Taxon labels C, G, Q, A and L refer to *Anopheles coluzzii*, *An. gambiae*, *An. quadriannulatus*, *An. arabiensis*, and *An. melas*, respectively, whereas Z is an outgroup. The arrow shows the hybridization event in the phylogenetic network, and the value associated with it (0.3) is the inheritance probability used. The values in black are node heights in coalescent units for the baseline setting (these values were scaled differently to produce different data sets; see main text). The node heights not displayed on the phylogenetic network are the same as their counterparts in the tree.

For each of the two species phylogenies in Figure 1, we generated data with three levels of ILS by scaling branch lengths in coalescent units by 1x (the values shown in Figure 1), 2x, and 5x, for all branches excepting the two immediately descending from the root. All scales used a branch length of 10 coalescent units for the branch leading to the ingroup in order to minimize ILS above the root node and ensure the validity of outgroup rooting, while avoiding saturation of sequence alignments. The branch leading to the outgroup taxon was adjusted to keep the phylogeny ultrametric.

We set population mutation rates *θ* to be 0.05, 0.025 and 0.01 for each scale respectively, to keep identical the branch lengths in units of substitutions per site for ingroup species and, consequently, keep the level of GTEE as uniform as possible across scales.

For each replicate, we used the program ms (Hudson, 2002) to simulate 10,000 conditionally independent gene trees for each exemplar species phylogeny. The number of replicates was 100 for each phylogeny. The following commands were used to generate gene trees on the species tree of Figure 1A with the 1x, 2x, and 5x branch length settings, respectively:

~~~
ms 6 10000 −T −I 6 1 1 1 1 1 1 −ej 0.15 6 5 −ej 0.2 3 2 −ej 0.4 5 4 −ej 0.5 4 2 −ej 5.5 2 1
ms 6 10000 −T −I 6 1 1 1 1 1 1 −ej 0.3 6 5 −ej 0.4 3 2 −ej 0.8 5 4 −ej 1.0 4 2 −ej 6.0 2 1
ms 6 10000 −T −I 6 1 1 1 1 1 1 −ej 0.75 6 5 −ej 1.0 3 2 −ej 2.0 5 4 −ej 2.5 4 2 −ej 7.5 2 1
~~~

The following commands were used to generate gene trees on the species network of Figure 1B with the 1x, 2x, and 5x branch length settings, respectively:

~~~
ms 6 10000 −T −I 6 1 1 1 1 1 1 −es 0.05 4 0.3 −ej 0.075 6 4 −ej 0.15 4 5 −ej 0.2 3 2
−ej 0.4 5 7 −ej 0.5 7 2 −ej 5.5 2 1
ms 6 10000 −T −I 6 1 1 1 1 1 1 −es 0.1 4 0.3 −ej 0.15 6 4 −ej 0.3 4 5 −ej 0.4 3 2
−ej 0.8 5 7 −ej 1.0 7 2 −ej 6.0 2 1
ms 6 10000 −T −I 6 1 1 1 1 1 1 −es 0.25 4 0.3 −ej 0.375 6 4 −ej 0.75 4 5 −ej 1.0 3 2
−ej 2.0 5 7 −ej 2.5 7 2 −ej 7.5 2 1
~~~

For each gene tree, we used the program `seq-gen` (Rambaut & Grass, 1997) to generate sequence alignments under a general time-reversible (GTR; Tavaré, 1986) model with base frequencies 0.2112, 0.2888, 0.2896, 0.2104 (A, C, G, T) and transition probabilities 0.2173, 0.9798, 0.2575, 0.1038, 1.0, 0.207 (A to C, A to G, A to T, C to G, C to T, T to G).

To generate data with heterogeneous rates across the *m* loci, we further sampled a vector of rates *M* = *μ*_1_, *μ*_2_,.., *μ*_*m*_ under the Dirichlet distribution Dir (*α*), where the concentration parameter *α* is a vector of *m* values all set to 1. Then we scaled the *i*-th gene tree by (*θ*/2) × *μ*_*i*_ as part of the `seq-gen` command to generate sequences.

To study the effect of GTEE on the methods, we generated sequences lengths for all loci using two different lengths: 2000 sites and 500 sites. The following command was used to generate homogeneous sequence data with substitution rate 1 for all loci for the 1x node height setting with *θ* = 0.05 and sequence length 500:

~~~
seq−gen −mGTR −s0.025 −f0.2112,0.2888,0.2896,0.2104
−r0.2173,0.9798,0.2575,0.1038,1.0,0.207 −l500
~~~

By changing the value after “-s” to (*θ*/2) × *μ*_*i*_ in the command, we generated sequences with locus-specific rates.

#### 3.1.1 Simulating smaller synthetic data sets

As full Bayesian inference of phylogenetic networks using the method `MCMC_SEQ` is computationally demanding for the larger number of taxa and loci, we restricted the model species phylogenies of Figure 1 to four ingroup taxa *An. coluzzii*, *An. quadriannulatus*, *An. arabiensis* and *An. melas*, in addition to the outgroup taxon. We limited this study to 2x and 5x scaled species branch lengths, and sequence alignment lengths of 2,000 sites. As above, we used the program `ms` to simulate 100 conditionally independent gene trees. The number of replicates was 10 for each of the two model phylogenies.

We generated two data sets for each replicate to compare the accuracy of phylogenetic network inference methods on data that have a single rate for all loci (the SR model, for “single rate”) as well as data with locus-specific rates (the DR model, for “Dirichlet rates”). We set the population mutation rate to *θ* = 0.01, and therefore the simulated gene trees were scaled by 0.005 to convert their branch lengths into expected substitutions per site.

### 3.2 Processing of *Heliconius* alignments

We reused the whole genome alignments of *Heliconius* species from Edelman et al. (2019). We extracted 10kb windows spaced at 50kb intervals using the `makewindows` command in `bedtools` v2.29.2. Then, we used *hal2maf* v2.1 to obtain alignments with reference genome *H. melpomene*. We converted the alignments from MAF to FASTA format with the tool `msa_view`. We then removed loci with a Jukes-Cantor distance between any pair of species greater than 0.2, as *Heliconius* butterflies are all closely related and such divergent loci may correspond to paralogs or pseudogenes. Due to the computational limits of full Bayesian inference of phylogenetic networks, we restricted the set taxa to three ingroup species, in addition to the outgroup species *Agraulis vanillae*, in two different ways: One set consists of ingroup species *H.erato.demophoon*, *H.hecalesia*, and *H.melpomene*, and the other consists of ingroup species *H.melpomene*, *H.timareta*, and *H.numata*. Finally, we filtered out loci with missing species, selected 100 loci at random, and truncated each locus to 500 sites to minimize intralocus recombination.

### 3.3 Species phylogeny and gene tree inference

For the simulated and butterfly data sets we estimated a gene tree for each locus independently of the species tree using `IQ-TREE` v1.6.10 (Nguyen et al., 2015), and rooted it at an outgroup. In the case of the simulated data, the outgroup was taxon Z. In the case of the biological butterfly data, the outgroup was *Agraulis vanillae*.

For the *Anopheles* data set, we reused the previously estimated bootstrap distributions of gene trees on autosomes (Wen, Yu, Hahn, et al., 2016) from the set of six species *An. coluzzii*, *An. gambiae*, *An. quadriannulatus*, *An. arabiensis*, *An. melas* and *An. merus*. These gene trees were inferred from previously published sequences (Fontaine et al., 2015). For each locus, we summarized the 100 bootstrap trees using a majority-rule consensus tree with a minimum clade frequency of 0.7, and resolved polytomies randomly.

Then we used `InferNetwork_ML` to infer species phylogenies from gene tree estimates. We ran the method with the number of runs set to 50 and the number of networks returned set to 10, with branch lengths and inheritance probabilities post-optimized, using the following command:

~~~
Infernetwork_ML (all) num-retic -po -di -x 50 -n 10;
~~~

where `num-retic` is an integer value that specifies the maximum number of reticulations allowed during the search for maximum likelihood estimates of the species phylogenies. When this value is set to 0, the method searches for a species tree.

For testing the impact of rate heterogeneity on species phylogeny inference, we set the maximum number of reticulations allowed to 0, 1, 2, and 3 to obtain the maximum likelihood estimates with the corresponding numbers of reticulations for both simulated and biological data.

For the full Bayesian analysis, we ran `MCMC_SEQ` on multi-locus alignment data and used the default priors to infer species phylogenies from the sequence data directly. For this analysis we assumed equal base frequencies and substitution rates, i.e. a Jukes-Cantor model of evolution (Jukes & Cantor, 1969). For the simulated data, we set chain lengths to 40,000,000, the burn-in period to 10,000,000, and the sample frequency to 1 in 5,000. The following command was used to sample species phylogenies with a single substitution rate:

~~~
MCMC_SEQ -cl 40000000 -bl 10000000 -sf 5000;
~~~

The following command was used to sample species phylogenies with varying substitution rates with DEO enabled by “-murate” (see Section 3.5 below):

~~~
MCMC_SEQ -cl 40000000 -bl 10000000 -sf 5000 -murate;
~~~

For the empirical butterfly data, we ran 10 chains with 80,000,000 iterations, a burn-in period of 8,000,000 iterations, and a 1-in-5,000 sampling frequency, allowing one population mutation rate per branch for both models of uniform substitution rates and variable substitution rates. We again assumed a Jukes-Cantor model of sequence evolution. Each chain was started with a unique random seed to ensure samples from the different chains were independent. We summarized the species network as the topology with the highest marginal probability from the samples across all chains after burn-in, and summarized the continuous parameters using their means and standard deviations. We summarized gene trees as the maximum clade credibility (MCC) tree with mean node heights.

### 3.4 To network or not to network: Test statistics

Many methods for the inference of species networks require pre-specifying the number of reticulations in the network. One statistical approach to estimating this number is to treat gene tree topologies as following a multinomial distribution, where the null hypothesis is that the sample of observed topologies is drawn from the distribution predicted by the multispecies (network) coalescent for the maximum likelihood species phylogeny with some number of reticulations. If the null hypothesis is rejected, this is interpreted as support for existence of additional reticulations. This approach is an application of the test introduced by Degnan and Rosenberg (2009).

While an exact statistical test is currently intractable for multinomial distributions with more than a few categories (Resin, 2022), several approximations exist. We evaluated three such approximations: Pearson’s *χ*^2^-test (Pearson, 1900), the *G*-test (Woolf, 1957), and a newer Monte Carlo-based approach (Resin, 2022). However, none of these tests directly account for GTEE which may also cause the null hypothesis to be rejected regardless of any introgression, a concerning possibility we have investigated by simulation.

Pearson’s *χ*^2^-test statistic is defined as

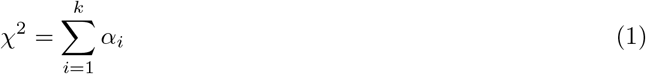

and the *G*-test is defined as

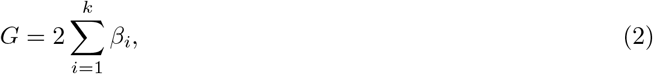

where the sums are taken over the *k* possible gene tree topologies given some number of taxa *n*. In the above equations,

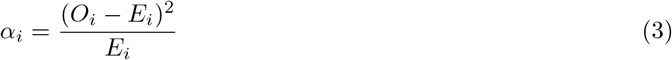

and

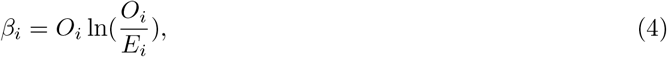

where *E*_*i*_ is the expected frequency of gene tree topology *i* under the null model, and *O*_*i*_ is its observed frequency. Both of these test statistics follow an approximately *χ*^2^ distribution when the null hypothesis is true (Larntz, 1978). The usual definition of the *G*-test appears undefined when *O*_*i*_ = 0, and while a number of approaches have been proposed to deal with zero counts (Hosmane, 1987), we chose to extend by continuity so that

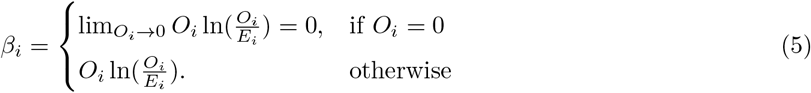

Under some conditions, e.g. larger values of *n* and/or lower levels of ILS, the expected frequencies of certain gene tree topologies will be close to zero, and the observed frequencies are typically zero or one. Pearson’s *χ*^2^ test is overly sensitive to these low expected frequencies, as *a*_*i*_ becomes very large when *E*_*i*_ is very small and *O*_*i*_ = 1, and the sum of *a*_*i*_ over all categories where *O*_*i*_ = 0 will be the sum of those expected frequencies, which may also inflate the test statistic. The *G*-test is more robust; when *O*_*i*_ = 1, *β*_*i*_ reduces to — ln(*E*_*i*_) which as a logarithm grows much more slowly than *α*_*i*_ as *E*_*i*_ approaches zero, and when *O*_*i*_ = 0, *β*_*i*_ does not contribute anything to the test statistic.

For the aforementioned test statistics *p*-values are straightforward to calculate, as both statistics approximate the *χ*^2^ distribution when the null hypothesis is true (Larntz, 1978). For five-taxon data sets, there were 105 unique rooted gene tree topologies (assuming only one sample per species) hence there were 105 − 1 − 3 = 101 degrees of freedom, as there were three internal branch lengths to be estimated by maximum likelihood. For three-taxon data sets, there are only three unique rooted gene tree topologies hence there was 3 − 1 − 1 = 1 degree of freedom, as there was only one internal branch length to be estimated by maximum likelihood.

We also included alternative approaches which do not use this approximation from the R package `ExactMultinom` v0.1.2 (Resin, 2022). For three-taxon data sets we used the exact multinomial test with the null hypothesis that the gene tree distribution follows a multinomial distribution defined by the MSC (Degnan & Salter, 2005). The exact multinomial test could not be used for these data sets because there were 105 gene tree topologies which exceeds the maximum of 15 categories permitted by the `ExactMultinom` implementation. Instead, we used the Monte-Carlo simulation based approach implemented in the same package when running the test on five-taxon data sets.

The above tests require knowledge of a species tree with branch lengths (in order to calculate gene tree distributions under the MSC) as well as gene tree topologies estimated on the individual loci. For each data set, we used the true gene trees, gene trees estimated from sequences of length 2000, and gene trees estimated from sequences of length 500 to reflect using correct gene trees, gene tree estimates with low levels of error, and gene tree estimates with high levels of error, respectively. For the species tree estimate, we used `InferNetwork_ML` on the set of gene trees as input while setting the maximum number of reticulations to zero. We used the command `CalGTProb` in `PhyloNet` to compute the gene tree topology probabilities.

We propose a new approach of applying the test statistics to three-taxon subsets of data sets, then aggregating the results. Here, the species tree on three taxa is estimated exactly by setting its topology to equal that of the topology of the gene tree with the highest probability (the “major gene tree topology”) and the length of its internal branch to

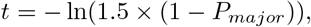

where *P*_*major*_ is the probability of the major gene tree topology.

Since we are now performing many tests for a single replicate, multiple-testing error correction must be used to avoid excessive false positives, and we evaluated the Bonferroni family-wise error-rate (FWER), the Simes–Hochberg FWER (Hochberg, 1988; Simes, 1986) and the Benjamini-Hochberg false discovery rate (FDR) methods (Benjamini & Hochberg, 1995) of error correction. We used the implementations in `statsmodels` v0.14.0 (Seabold & Perktold, 2010).

We also ran two tests developed specifically for testing for the presence of introgression in the evolutionary history of a given genomic data set. The test of Cai and Ané (2021) evaluates the fitness of the MSC or MSNC based on quartets given a candidate species phylogeny and a set of observed gene tree frequencies obtained from multiple loci, and can be applied to data sets with any number of taxa. The *D*_3_ test of Hahn and Hibbins (2019) is applicable to data sets with three taxa. Assuming a species tree ((A,B),C), the test is defined by

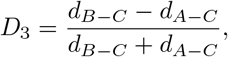

where *d* denotes pairwise distances between taxa using the multi-locus data, and a *p*-value is computed from a z-distribution of *D*_3_ values computed via bootstrapping.

### 3.5 Accounting for rate heterogeneity in MCMC_SEQ

Instead of assuming a common substitution rate for all loci, we now incorporate a vector of substitution rates *M* = *μ*_1_, *μ*_2_,.., *μ*_*m*_ in our model, where *m* is the number of loci. Using a multiplier *δ* and a vector of weights for the locus-specific substitution rates *W* = *w*_1_, *w*_2_,.., *w*_*m*_, the DEO proposes changing two rates in *M* at a time. The first step is to select two indices of substitution rates *d*_1_ and *d*_2_ with weights 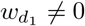 and 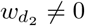. Then, the operator computes a value *x* = *r* × *δ*, where *r* is a random number from the unit range. Finally, we have 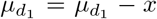, and 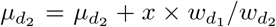 as the proposed values for the substitution rates. Note that this proposal imposes a flat Dirichlet prior on the element-wise product of *M* and *W* and holds the sum of this product constant, ensuring the mean substitution rate is unchanged (Figure 2).

**Figure 2:**
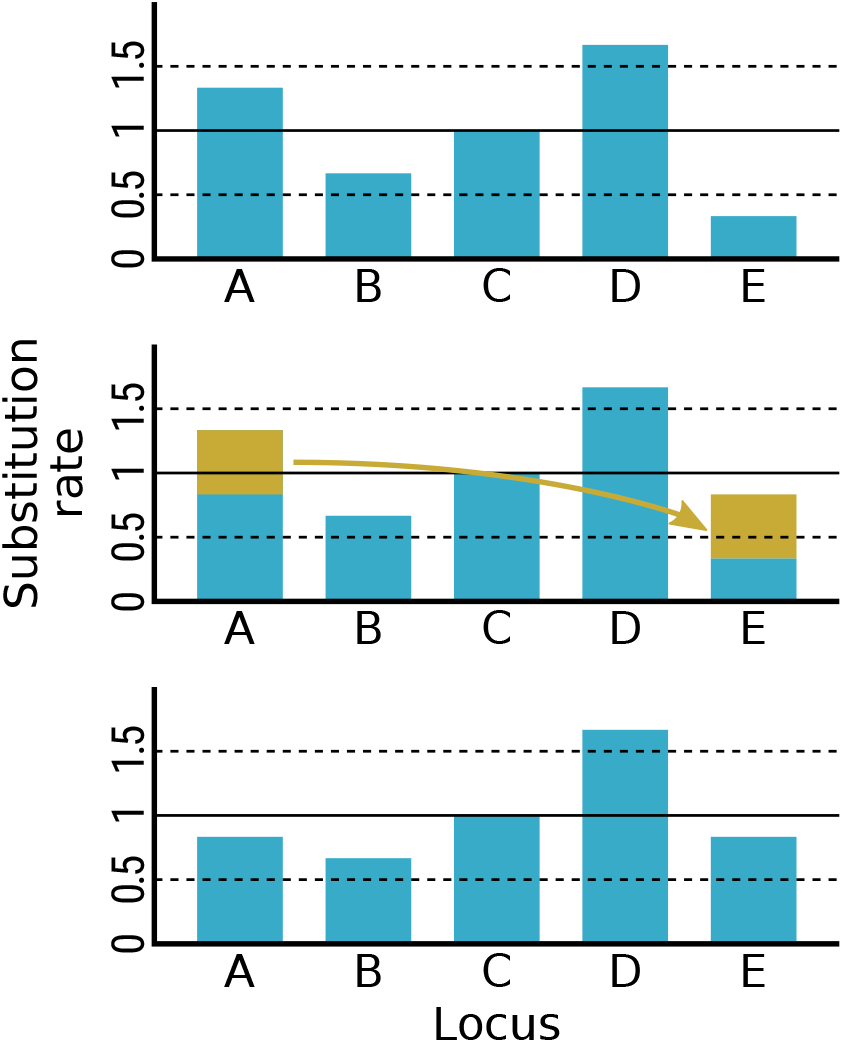
Illustration of the Delta Exchange Operator (DEO). The current state has a mean substitution rate of 1 (solid horizontal line, top). The substitution rate of locus A is proposed to be decreased and the rate of locus E is proposed to be increased by an equal amount (amounts and transfer shown in gold, middle). If accepted, this proposal maintains the mean substitution rate of 1 (solid horizontal line, bottom).

### 3.6 Quantitative measures of accuracy and error

We measured the topological accuracy of phylogenetic network estimates using the distance metric of Nakhleh (2009), henceforth referred to as network distance. We measured the accuracy of gene tree estimates using the normalized Robinson-Foulds measure for topological distance (nrRF; Robinson & Foulds, 1981) and the normalized branch score measure for topology and branch length distance (nrBS; Heled & Drummond, 2009).

The nrRF distance between two trees *T* and 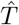, which in our case would be the true and estimated trees, respectively, is defined as the number of clades present in only one of the two trees divided by the total number of distinct clades in the two trees.

The nrBS is defined as

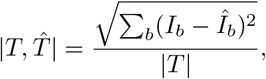

where the sum is taken over all branches in the true tree *T*, *I*_*b*_ is the length of branch *b* in the true tree *T*, and 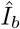 is length of branch *b* in the estimated tree 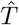.

For assessing the accuracy of substitution rate estimates we used the relative error of substitution rate at locus *i* calculated as 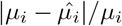, where *μ*_*i*_ is the true substitution rate and 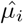 is the estimated substitution rate. The average relative error over all loci was calculated as 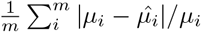.

## 4 Results

### 4.1 Statistical tests for reticulate evolution

As mentioned in Section 3.4, we varied the level of GTEE by using the original simulated gene trees (no GTEE), using gene trees inferred from 2000-site simulated sequence alignments (low GTEE), and gene trees inferred from 500-site simulated sequence alignments (high GTEE). By varying the sequence alignment length, the topological error in estimated gene trees was effectively spread out, as the median gene tree inferred from 2000-site alignments had more than double the topological error of the median gene tree inferred from 500-site alignments (Figure 3).

**Figure 3:**
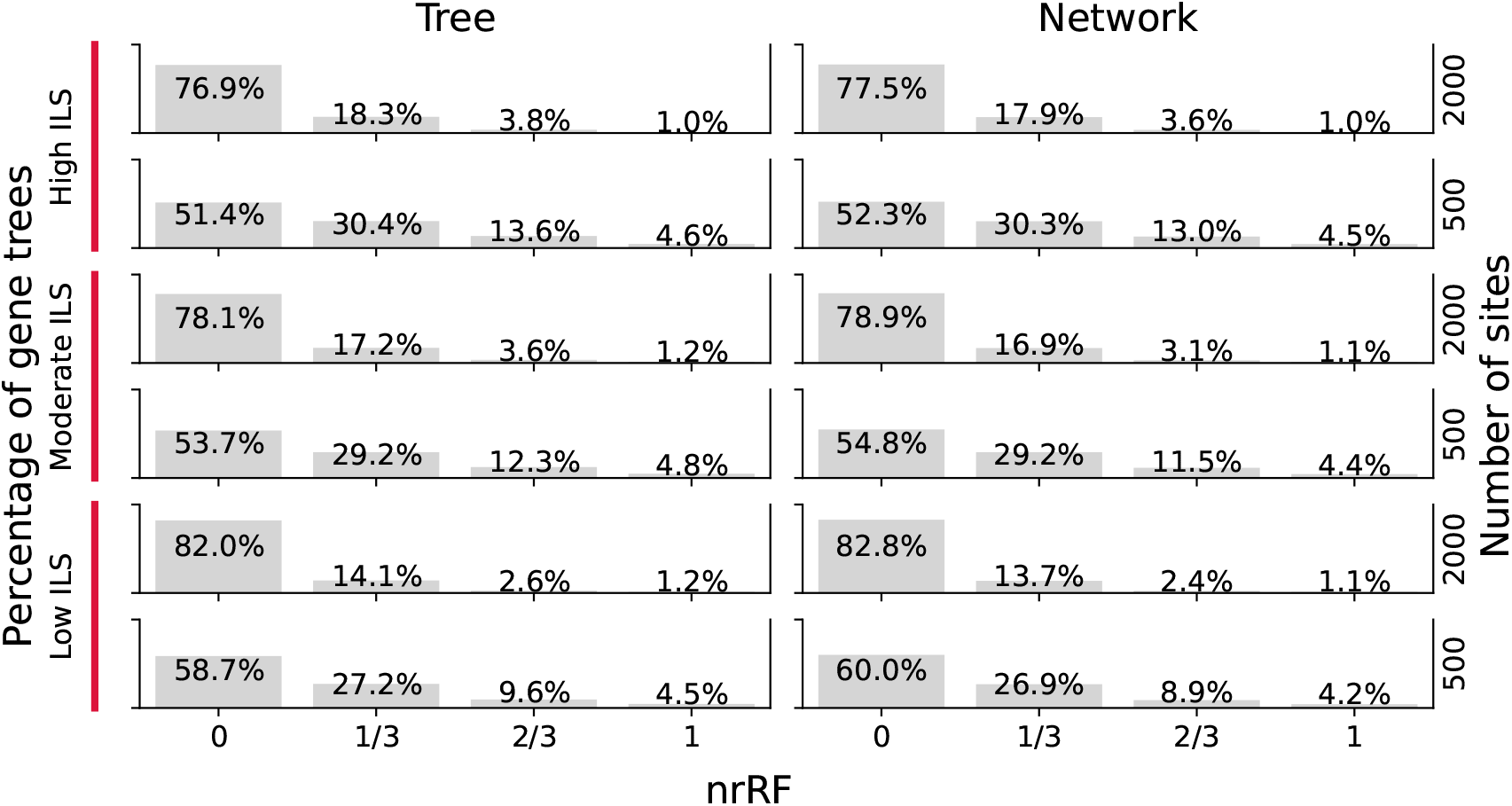
Maximum likelihood gene tree estimation error. Normalized Robinson-Foulds distances (nrRF) were calculated for pairs of simulated and maximum likelihood estimated gene trees. There were only four possible values of nrRF, proportional and corresponding to the number of correct clades, besides the root and crown of the ingroup, present in the inferred gene tree. Besides nrRF distance, gene trees were grouped by whether they were simulated under a species tree or network, the number of sites in the alignment used to infer the gene tree, and the level of incomplete lineage sorting (ILS) as determined by the scale of the species phylogeny.

#### 4.1.1 Existing statistical approaches to detect reticulations have high false positive rates

When data were simulated following a five-taxon species tree without any reticulations, and the maximum likelihood species tree without reticulations was used as the null hypothesis, all rejections of the null hypothesis must be false positive indications of a reticulate evolutionary history. Using a significance threshold of *p* = 0.05 we expect 5% of tests to be rejected for a false positive rate of 5%, which we observed for all three approximate multinomial tests in the absence of GTEE when ILS was high (Figure 4).

**Figure 4:**
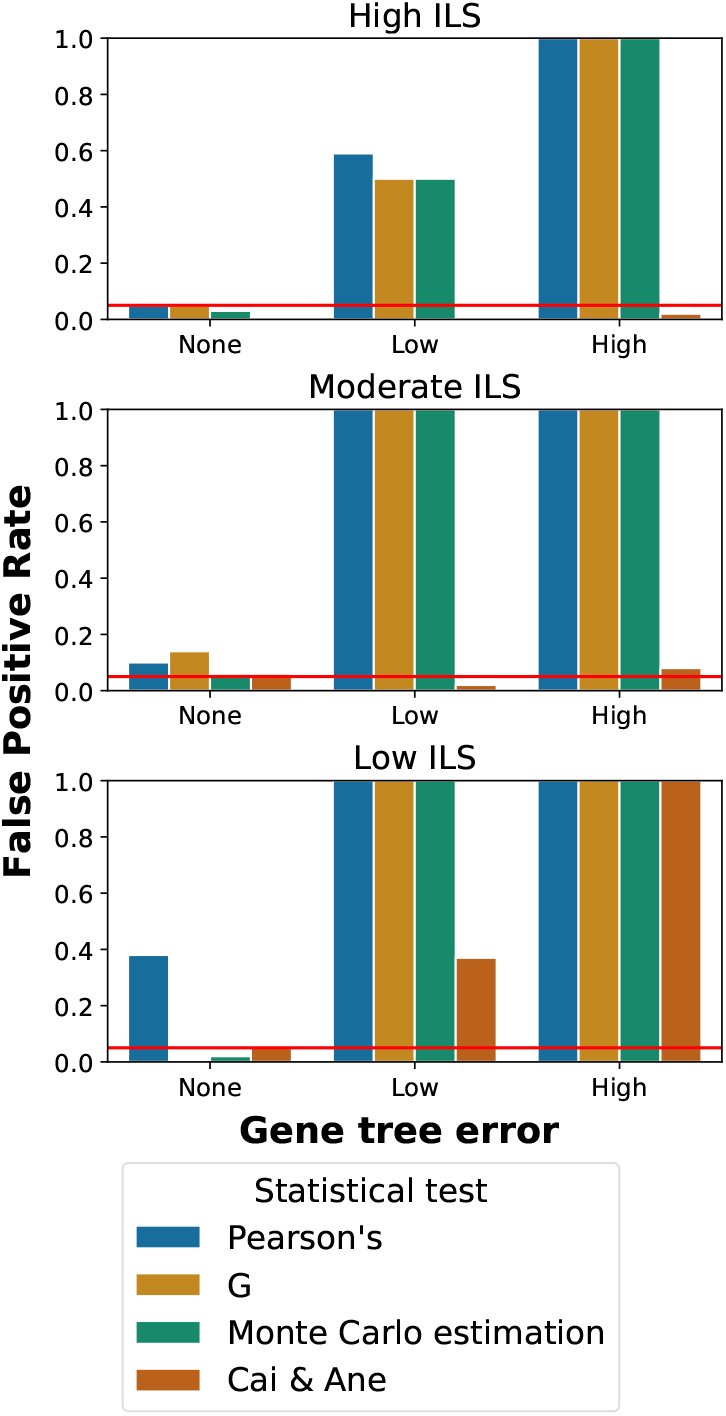
The false positive rate of existing statistical approaches of testing for reticulations when the true species phylogeny is a tree. The false positive rate corresponds to the proportion of tests rejected with a *p*-value below the statistical threshold of 0.05 (red horizontal line). Incomplete lineage sorting (ILS) was varied by changing the branch lengths of the species tree in coalescent units (but not substitutions per site), and gene tree estimation error was varied by using the actual simulated trees for no error, trees inferred from long (2000 site) alignments for low error, and trees inferred from short (500 site) alignments for high error.

However, when the level of ILS is decreased, the number of unobserved gene tree topologies will increase for a fixed sample size, leading to excessively large Pearson’s test statistics. This is a known problem of Pearson’s *χ*^2^-test when probability distributions are highly skewed, as gene tree topology distributions will be when ILS is low (Bradley et al., 1979). As a result, the false positive rate of Pearson’s test was excessively high under the low ILS regime. Both the *G*-test and the Monte Carlo approximation of the exact test fared better than those other approaches in the absence of GTEE, with a false positive rate of around or below 5% (Figure 4).

However, once even a low level of GTEE was introduced, all three methods rejected far more than 5% of species trees. When ILS was moderate or high, or when GTEE was high, all species trees were (inaccurately) rejected (Figure 4).

An alternative to this simple approach is that of Cai and Ané (2021), which builds on the TICR test (Stenz et al., 2015). These more complex approaches extract quartet concordance factors from a set of sampled gene tree topologies, calculate *p*-values for those quartet concordance factors, then compare outlier *p*-values to an expected distribution. Because a single gene tree may contribute to multiple quartets, these *p*-values are not independent, and Cai and Ané attempt to correct for this dependence using simulation. Their approach is far more robust than approximate multinomial tests of gene tree topology distributions for five taxa, with false positive rates of around or under 5% except where ILS was low and GTEE existed (Figure 4). When ILS is low, sample sizes of discordant topologies will be small and hence the relative difference between expected and observed values caused by GTEE will be magnified, potentially explaining the poor performance in this case.

All four statistical tests rejected the null hypothesis for every replicate when the truth was a network (Figure 5), indicating that none of these methods suffer from low sensitivity under any tested condition.

**Figure 5:**
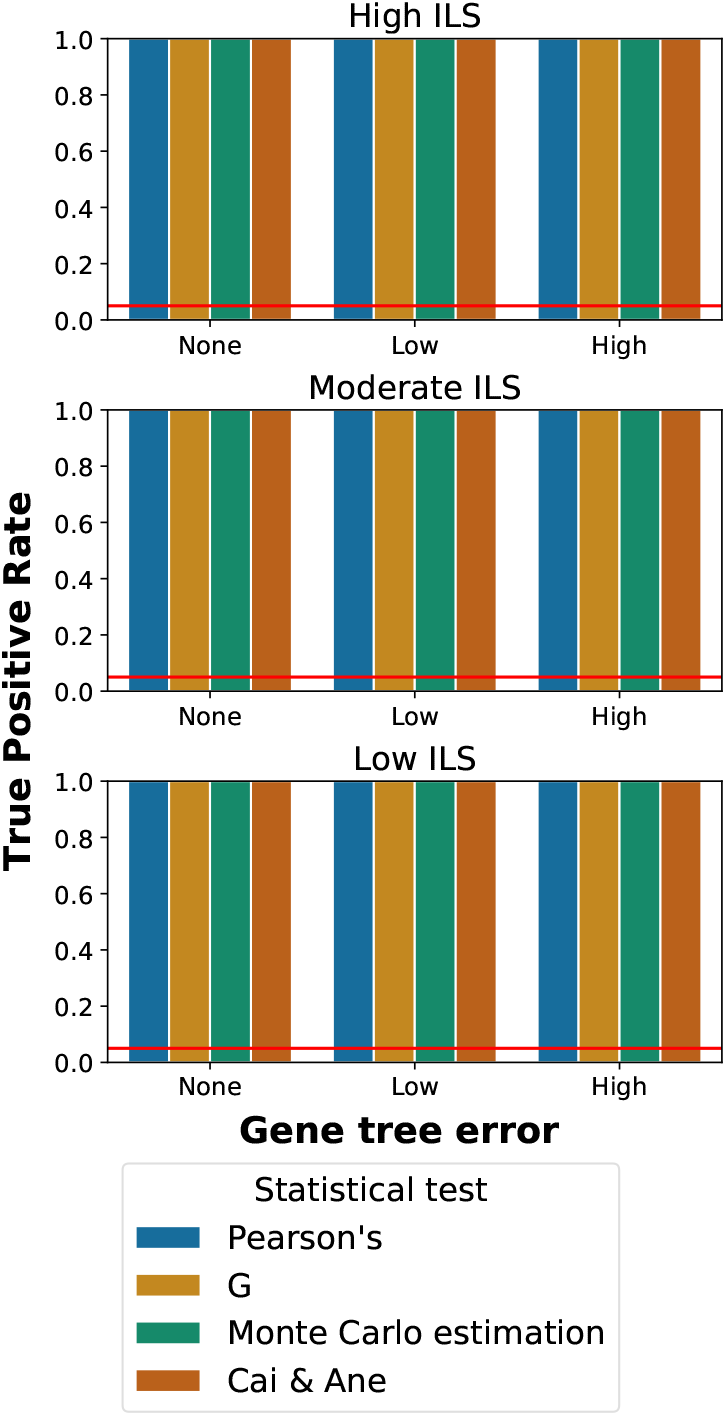
The true positive rate of existing statistical approaches of testing for reticulations when the true species phylogeny is a network. The true positive rate corresponds to the proportion of tests rejected with a *p*-value below the statistical threshold of 0.05 (red horizontal line). Incomplete lineage sorting (ILS) was varied by changing the branch lengths of the species tree in coalescent units (but not substitutions per site), and gene tree estimation error was varied by using the actual simulated trees for no error, trees inferred from long (2000 site) alignments for low error, and trees inferred from short (500 site) alignments for high error.

Each statistical test was performed on a server equipped with 2.2GHz Intel Xeon Gold 5220R CPUs, hence the absolute time required for each test is specific to that system. Regardless of the statistical method used, most of the running time was spent to estimate gene trees and the species phylogeny, which took on average 37.52 and 78.42 minutes per replicate. The multinomial tests all finished within 1 second for each replicate, while the method of Cai & Ané took on average 7.15 minutes.

#### 4.1.2 Calculating test statistics on triplets of taxa overcomes gene tree estimation error

Since all existing methods we evaluated that could test for reticulations in five-taxon species phylogenies were unreliable, we explored the applicability of a divide-and-conquer approach based on testing all three-taxon subtrees or subnetworks of the complete species phylogeny. Given the problems with the existing methods may in part stem from low sample sizes of unique gene tree topologies, rationally this approach should alleviate the problem given the number of unique topologies for each triplet is only three. Since the number of gene trees remains the same, the average number of observations of each unique topology will be substantially increased.

This divide-and-conquer approach also has additional computational advantages when testing a species tree. First, maximum likelihood inference of the internal branch length can be solved following a simple equation (Degnan & Rosenberg, 2009) rather than using a heuristic algorithm such as hill-climbing. Second, with only three categories, an exact multinomial test may be used in place of approximations (Resin, 2022). By testing species triplets instead of entire trees, we can also evaluate the use of the *D*_3_ test statistic that was developed to test for introgression using sequence alignments on three taxa (Hahn & Hibbins, 2019).

For the gene tree topology approaches, the null hypothesis is the same as before, that the sampled topologies are being drawn from the distribution predicted by the MSC. For *D*_3_, the null hypothesis is that the genetic distance from the most distantly related taxon to one of the two closely related taxa will be the same as the distance from the most distant to the other of the two closest taxa (Hahn & Hibbins, 2019). In either case, we interpret the rejection of the null hypothesis as indicating the presence of reticulation.

When the null hypothesis for one or more triplets is rejected, we interpret that as rejecting the null hypothesis for the whole species phylogeny. As discussed above, we evaluated the Bonferroni family-wise error-rate (FWER), the Simes–Hochberg FWER, and the Benjamini-Hochberg false discovery rate (FDR) methods for multiple-testing error correction.

All three multinomial tests—Pearson’s *χ*^2^ test, the *G*-test and the exact test—had a false positive rate of around or below 5% when the true species phylogeny is a tree (Figure 6).

**Figure 6:**
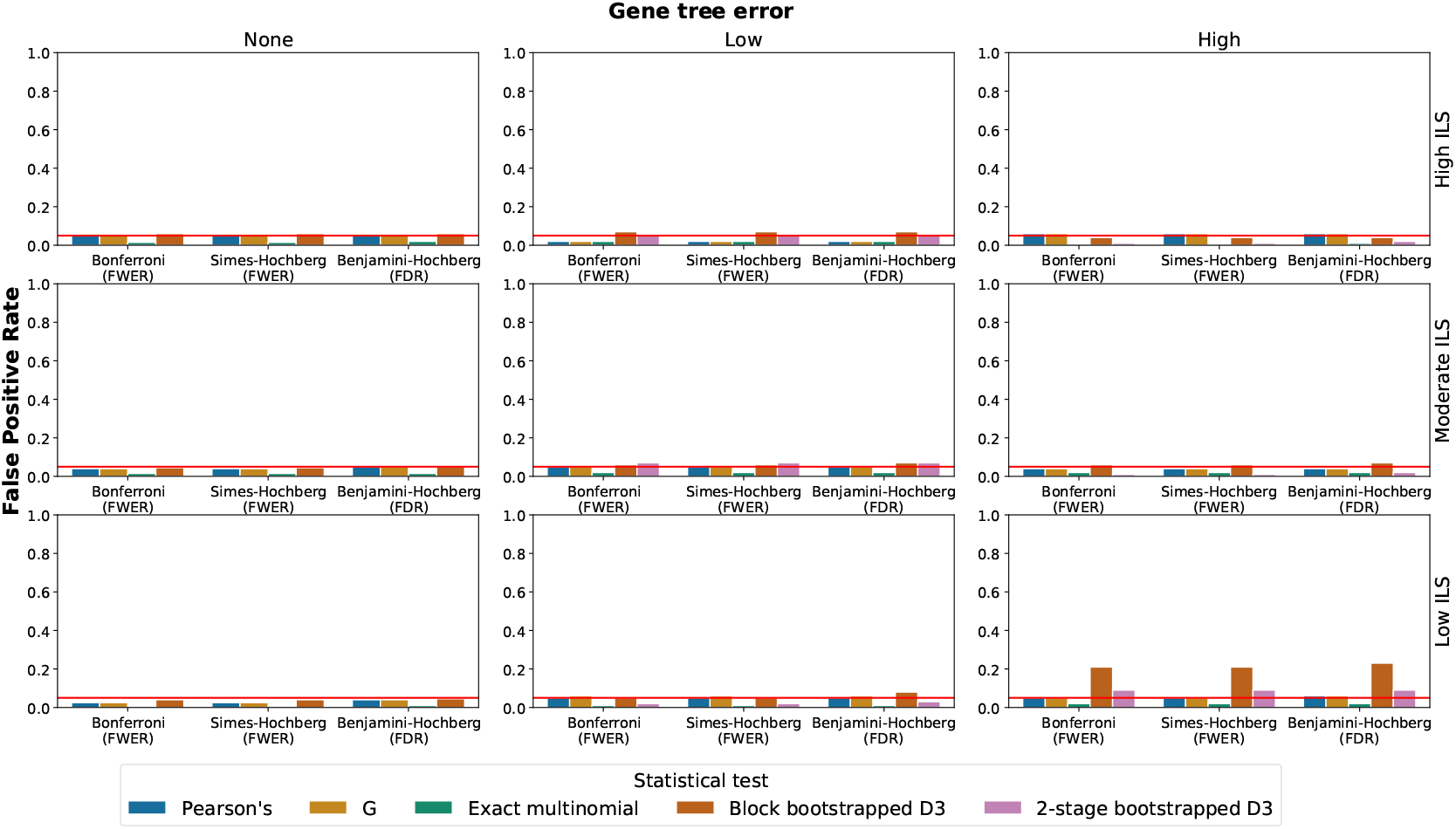
The false positive rate of divide-and-conquer approaches of testing for reticulations when the true species phylogeny is a tree. The false positive rate corresponds to the proportion of tests rejected with a *p*-value below the statistical threshold of 0.05 (red horizontal line). Incomplete lineage sorting (ILS) was varied by changing the branch lengths of the species tree in coalescent units (but not substitutions per site), and gene tree estimation error was varied by using the actual simulated trees for no error, trees inferred from long (2000 site) alignments for low error, and trees inferred from short (500 site) alignments for high error. Results are grouped by the family-wise error rate (FWER) or false discovery rate (FDR) method of multiple testing correction which was used.

There was minimal loss of sensitivity, with the null hypothesis of tree-like evolution rejected for all replicates where genes evolved following a species network with reticulation (Figure 7). These results were robust to the choice of multiple-testing correction method, to the degree of GTEE, and also to the level of ILS. This divide-and-conquer approach of testing triplets of species instead of the entire set of taxa therefore solves the problem of excessive false positives under the range of settings we studied.

**Figure 7:**
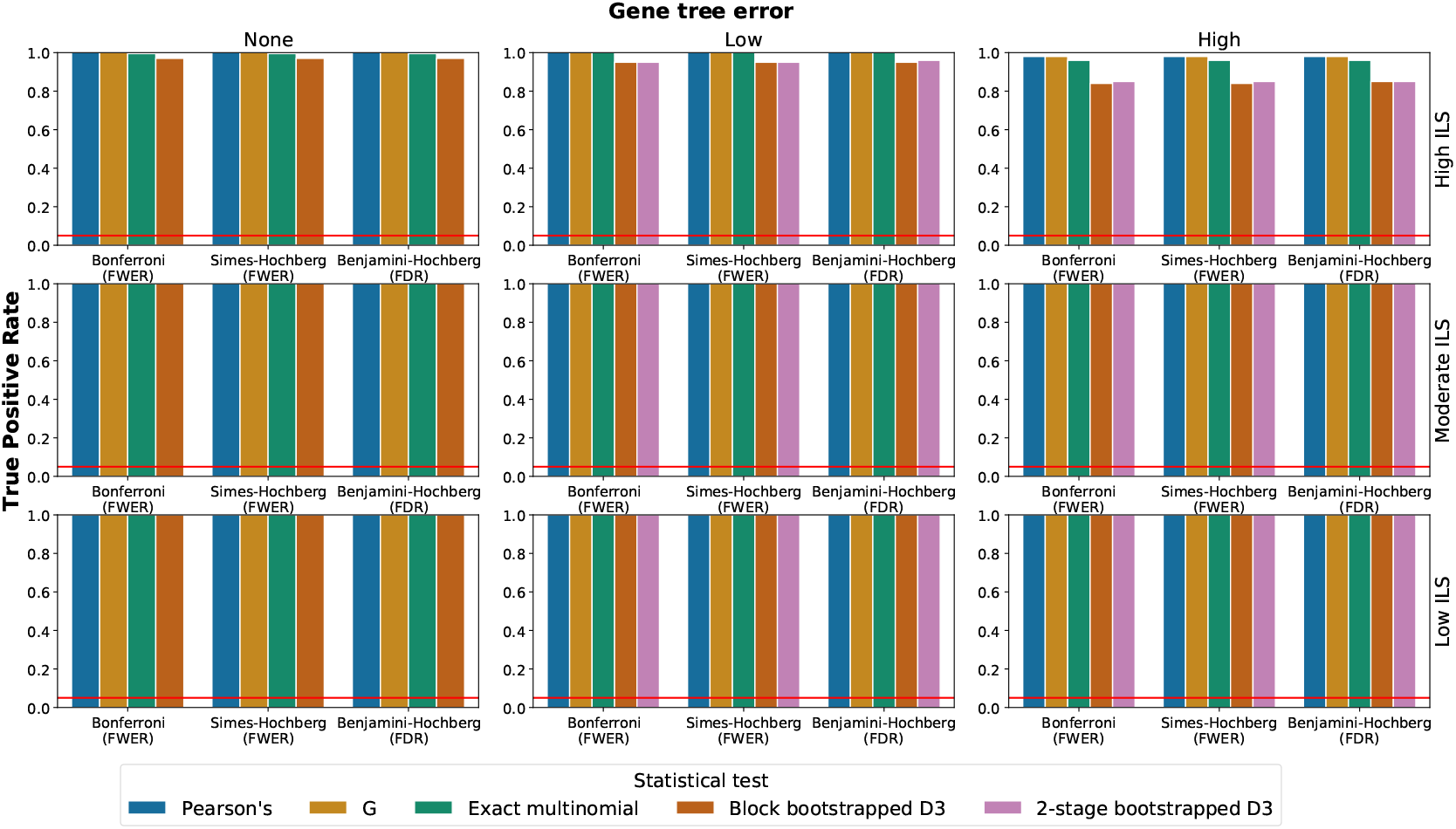
The true positive rate of divide-and-conquer approaches of testing for reticulations when the true species phylogeny is a network. The false positive rate corresponds to the proportion of tests rejected with a *p*-value below the statistical threshold of 0.05 (red horizontal line). Incomplete lineage sorting (ILS) was varied by changing the branch lengths of the species tree in coalescent units (but not substitutions per site), and gene tree estimation error was varied by using the actual simulated trees for no error, trees inferred from long (2000 site) alignments for low error, and trees inferred from short (500 site) alignments for high error. Results are grouped by the family-wise error rate (FWER) or false discovery rate (FDR) method of multiple testing correction which was used.

The original *D*_3_ test was described using moving block-bootstrapping to estimate statistical significance (Hahn & Hibbins, 2019). Expected distribution of *p*-values calculated using this bootstrap method differ from the those calculated directly from the data, so to avoid this problem, we instead applied the non-overlapping block-bootstrap (Härdle et al., 2003). Unlike for the multinomial tests, when using block-bootstrapping to test for tree-like evolution using the *D*_3_ statistic, we observed an excessive false positive rate when GTEE was high and ILS was low (Figure 6).

Since this excessive rate was not observed when the true genetic distances were used to calculate *D*_3_ (Figure 6), we hypothesized that this was caused by the failure of block-bootstrapping to account for the sampling error within each block. To account for this sampling error we applied 2-stage bootstrapping (Seo, 2008) which ameliorated the false positive rate. However, the sensitivity of the *D*_3_ test was lower than multinomial tests when ILS was high and GTEE existed regardless of which bootstrapping method was used (Figure 7).

The multinomial tests on triplets of species required estimating gene trees, which were reused from the monolithic approach. The test statistics themselves took less than one second per replicate. While, unlike the multinomial methods, *D*_3_ does not require estimated gene trees, bootstrapping of sequence alignments still took a substantial amount of time; testing for tree-ness took 10.09 minutes per replicate on average with block-bootstrapping, or 18.45 minutes using 2-stage bootstrapping.

### 4.2 Rate heterogeneity and species network inference

When GTEE results from the stochastic nature of molecular evolution and the limited information available in each locus, this error should not cause full Bayesian multilocus methods to infer spurious reticulations. This is because these methods jointly infer gene trees with the species phylogeny, integrating over all possible gene trees at each locus, unlike the statistical tests which (except for *D*_3_) all treat point estimates of gene trees as data regardless of the level of error in those point estimates. However, when GTEE is caused by model misspecification, these methods may no longer be robust. One form of model misspecification may be assuming all sequences evolved for all time under a single rate (SR), instead of each locus evolving at a different rate (DR).

Previous versions of the full Bayesian multilocus method `MCMC_SEQ` only implemented an SR model, so to study the effect of this misspecification we implemented a DR model of sequence evolution using the DEO as it is implemented in BEAST (Drummond & Rambaut, 2007). As in the study of statistical tests, we used a simulation approach, reusing the same species phylogenies (Figure 1), but with leaf G removed and the number of gene trees set to be 100 to reduce the time needed for `MCMC_SEQ` to converge. Gene trees were simulated under low and moderate ILS conditions, and sequence alignments simulated under SR or DR conditions were 2000 sites in length.

#### 4.2.1 Not accounting for rate heterogeneity causes full Bayesian methods to infer spurious reticulations

We first looked at the effects of rate heterogeneity under low ILS conditions. When data were simulated under the SR model and lacked rate heterogeneity, using the DEO implementation of the DR model for inference did not have a substantial negative effect on topological accuracy. However, when data were simulated under the DR model, and the SR model was used for inference by disabling the DEO, we observed a large decrease in accuracy (Figure 8, top row).

**Figure 8:**
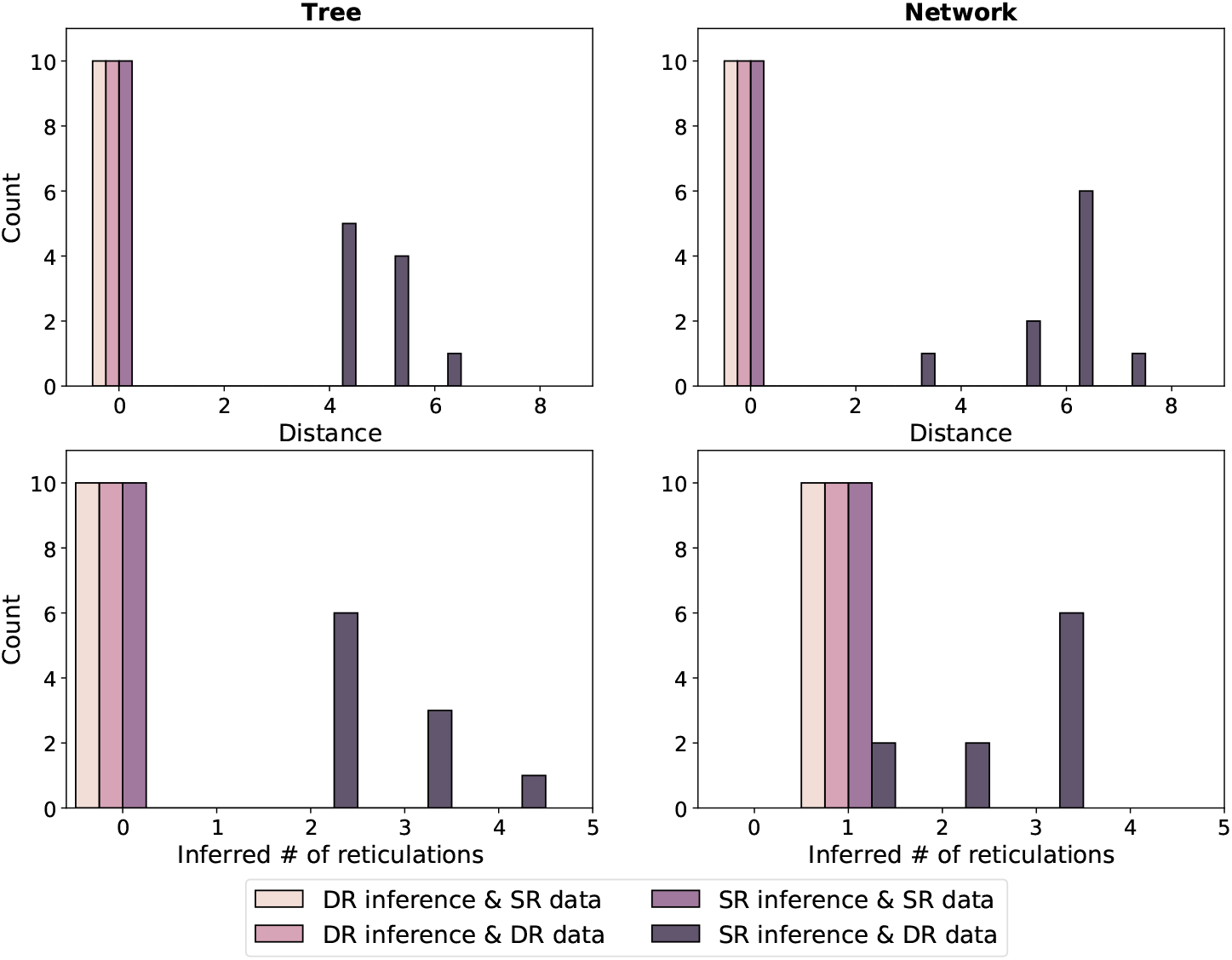
The effect of per-locus substitution rate variation on the accuracy of species phylogenies inferred using a full Bayesian approach. `MCMC_SEQ` was used to infer the species phylogeny together with the gene trees from multiple sequence alignments in a full Bayesian approach, under either single rate (SR) or Dirichlet rates (DR) models of per-locus substitution rate variation. Sequence alignments were also simulated under either SR or DR models. Each bar represents the number of replicates a given network distance from the true species phylogeny, for the exemplar species tree and network.

Further investigation revealed that the basis for this error was the inference of reticulations beyond the true number of reticulations (Figure 8, bottom row). Similar results were observed for the species network with the highest probability marginalized over branch lengths and other continuous parameters (Figure S1).

Under moderate ILS conditions the results were similar (Figures S2, S3), although the inferred species networks were sometimes inaccurate even without model misspecification. This may have been due to the lower correlation between gene and species phylogenies making individual loci less informative regarding the species phylogeny.

### 4.2.2 Summary methods are robust to substitution rate heterogeneity

Maximum likelihood methods of species network inference, unlike Bayesian methods, are not inherently able to estimate the number of reticulations present. This is because a network of higher likelihood should always be possible to find by increasing the number of reticulations *r*, even beyond the true *r* (Cao et al., 2023). Therefore a maximum likelihood search should return a network where *r* is equal to the maximum permissible value *r*_*max*_ set by the user or implementation. When *r*_*max*_ is set equal to the true *r*, introducing rate heterogeneity has only a very minor negative effect on species network accuracy when gene trees are estimated (Figure 9), despite a more substantial decrease in the accuracy of inferred gene tree topologies (Figure 10A). When the true phylogeny is a network, using estimated gene trees has a very negative effect on accuracy (Figure 9), but adding rate heterogeneity does not make it any worse.

**Figure 9:**
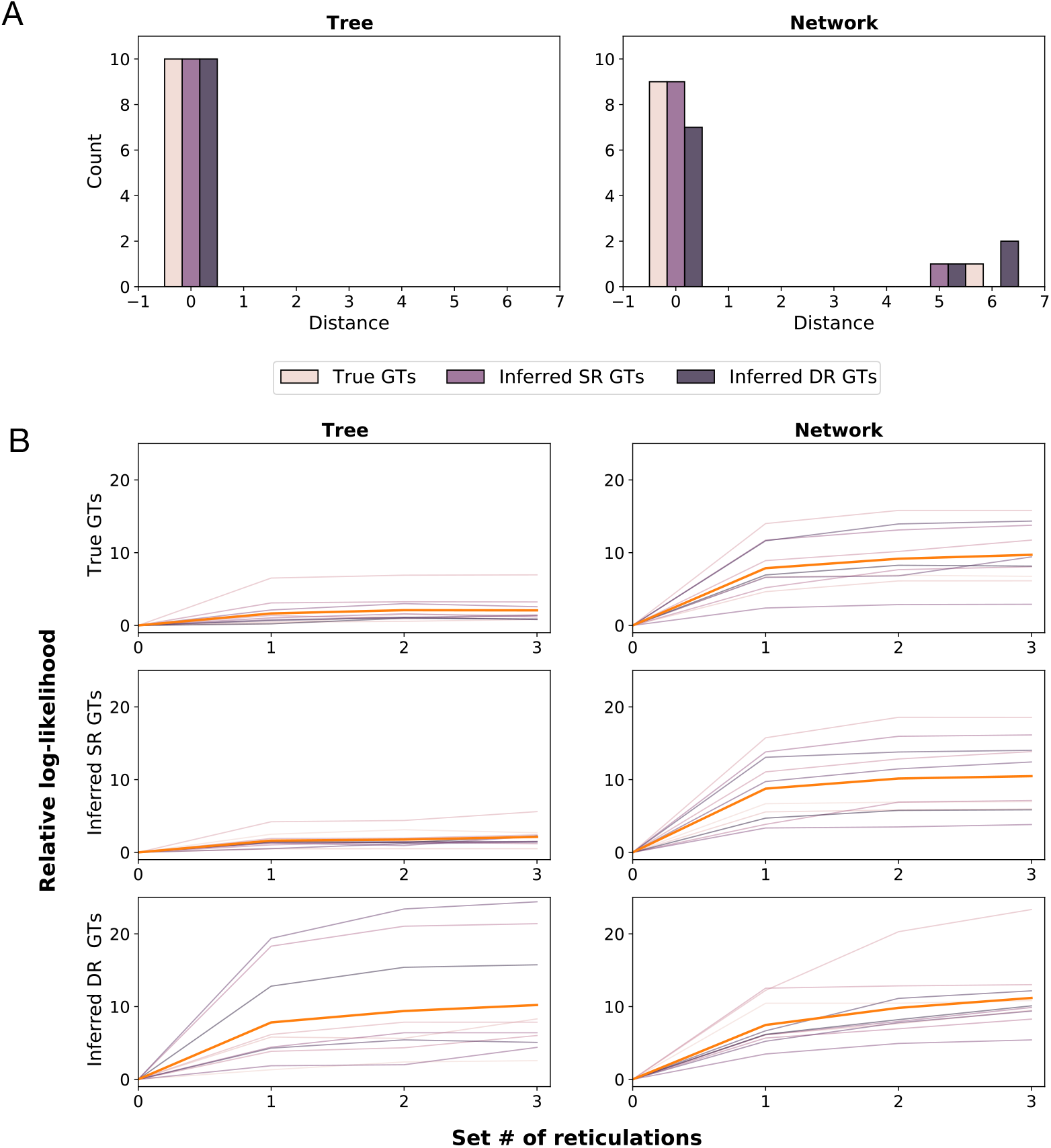
The effect of per-locus substitution rate variation on the accuracy of species phylogenies inferred by the maximum likelihood summary method `InferNetworks_ML`. (A) Each bar represents the number of replicates with a given network distance from the true species network for the two exemplar species phylogenies (a species tree and a species network with one reticulation). Gene trees used as input were either the true gene trees (True GTs), or inferred using `IQ-TREE` from sequence data simulated along those trees under a single rate model (inferred SR GTs), or a Dirichlet rates model (inferred DR GTs). (B) Log-likelihood increase of species networks identified using the maximum likelihood summary method `InferNetworks_ML`. The log-likelihood for a given maximum number of reticulations is relative to the log-likelihood of the zero-reticulation maximum likelihood network (i.e. the species tree). Gene trees used as input were either the true gene trees (True GTs), or inferred using `IQ-TREE` from sequence data simulated along those trees under a single rate model (inferred SR GTs), or a Dirichlet rates model (inferred DR GTs). Thick orange lines show the average increase relative to zero reticulations, all other lines show the increase for each individual replicate.

**Figure 10:**
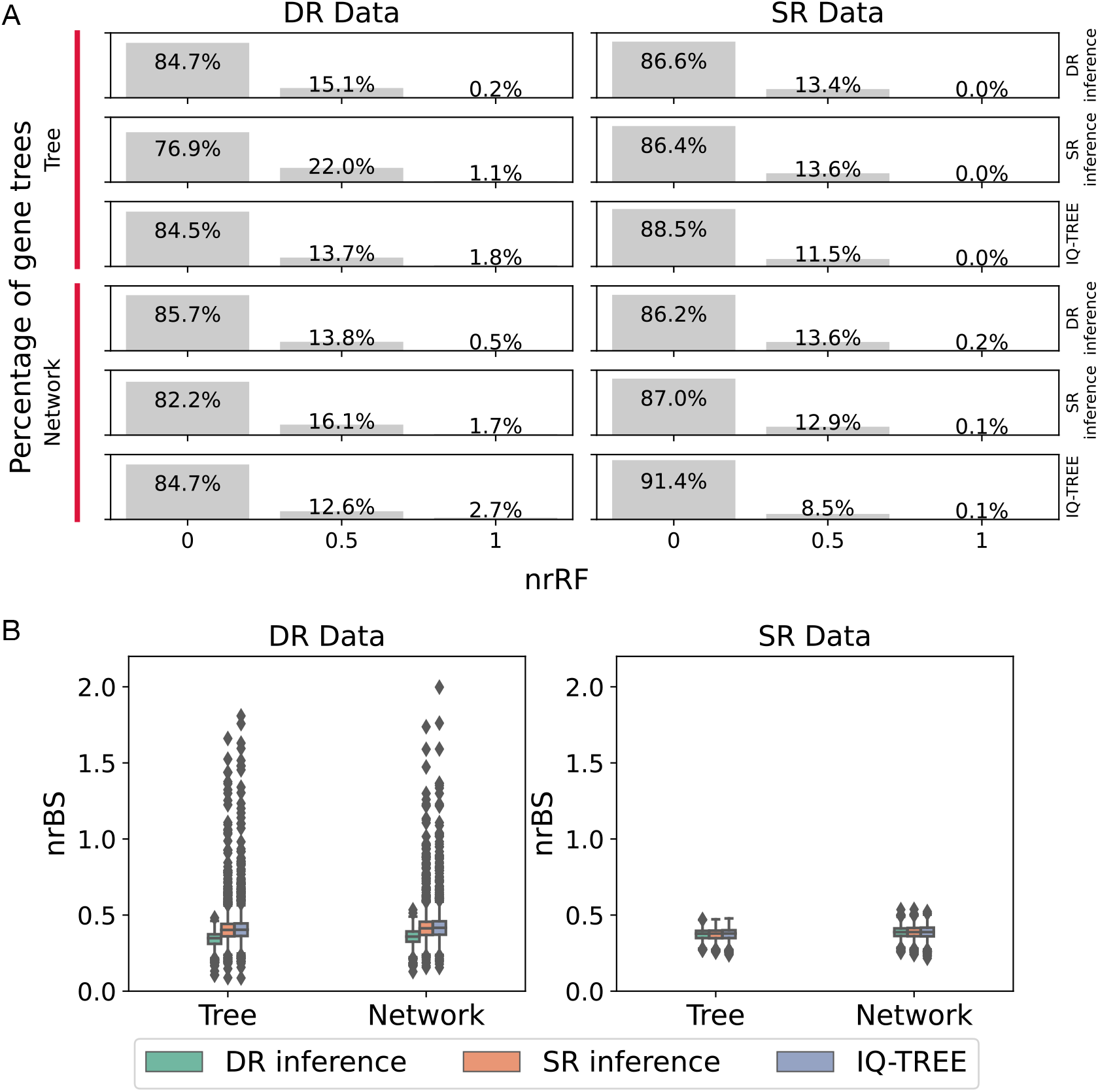
The effect of per-locus substitution rate variation on the accuracy of gene trees inferred independently or using a full Bayesian approach. (A) The accuracy of inferred gene tree topologies measured by normalized Robinson-Foulds (nrRF) distances. There were only three possible values of nrRF, proportional and corresponding to the number of correct clades, besides the root and crown of the ingroup, present in the inferred gene tree. (B) The accuracy of inferred gene trees including their branch lengths measured by normalized branch score (nrBS). `IQ-TREE` was used to infer gene trees independently of each other or the species phylogeny. `MCMC_SEQ` was used to infer posterior distributions of gene trees together with the species phylogeny from multiple sequence alignments in a full Bayesian approach, under either single rate (SR) or Dirichlet rates (DR) models of per-locus substitution rate variation, which were summarized as maximum clade credibility (MCC) point estimate trees with mean node heights before calculating nrRF and nrBS values. Sequence alignment data were simulated under either SR (right boxes) or DR (left boxes) models.

The noticeable difference between the accuracy of species networks and trees inferred from estimated gene trees cannot be attributed to differences in GTEE, as gene trees estimated using `IQ-TREE` were actually slightly more accurate when the true species phylogeny was a network (Figure 10A). Adding rate heterogeneity to the true gene trees has no effect since this summary method ignores branch lengths, and the true gene tree topologies are unaffected by rate heterogeneity.

The original publication on maximum likelihood species network inference (Yu et al., 2014) suggested increasing *r* until the increase in likelihood becomes negligible. This creates the appearance of a “shoulder” when the maximum likelihood is drawn as a function of *r*. The number of reticulations is able to be identified when the phylogeny is inferred with true gene trees or gene trees estimated under SR data, since we observe little growth in the likelihood after the point of the true number of reticulations (Figure 9, top and middle rows). When using DR data, the turning point becomes misleading for the species tree (Figure 9B, bottom left).

#### 4.2.3 Gene tree estimation

When data were simulated without rate heterogeneity under the SR model, `IQ-TREE` performed slightly better than `MCMC_SEQ` when inferring gene tree topologies regardless of whether the DEO implementation of the DR model was enabled (Figure 10A, right). While joint inference of species and gene phylogenies is typically superior to independent gene tree inference (Szöllősi et al., 2014), in this case IQ-TREE had the advantage of being able to estimate the GTR model parameters used to simulate sequence alignment, whereas `MCMC_SEQ` assumed equal base frequencies and substitution rates.

When data were simulated under the DR model with rate heterogeneity, using the DR model for inference with `MCMC_SEQ` outperformed both the SR model and `IQ-TREE` (Figure 10A, left). When considering branch lengths in addition to topology, the accuracy of all three methods was virtually indistinguishable when data were simulated without rate heterogeneity, but in the presence of rate heterogeneity enabling the DR model was clearly superior (Figure 10B).

The relative substitution rates estimated under the DR model using the DEO implementation in `MCMC_SEQ` were strongly correlated with the true simulated rates (Figure 11), although the average relative error was 18.81%, reflecting the limited information present in each individual sequence alignment. Estimation of relative rates is not possible using `IQ-TREE` due to the confounding of rates and time.

**Figure 11:**
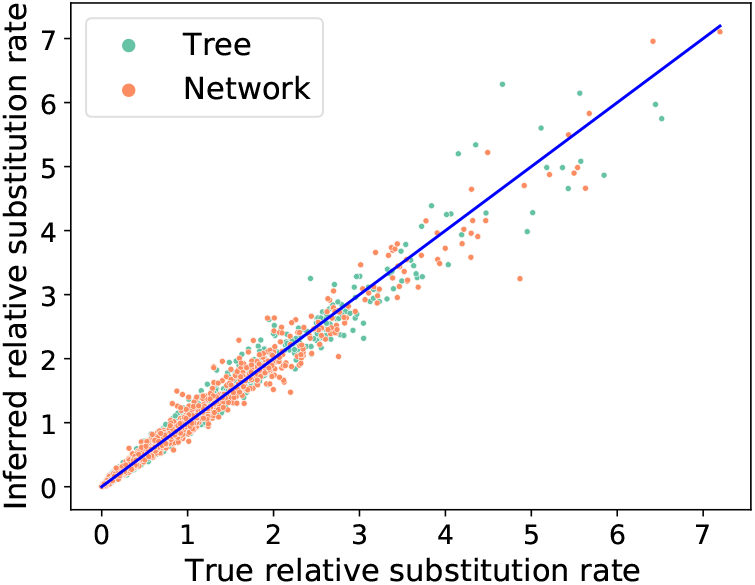
Per-locus substitution rate estimates using the Dirichlet rates model of per-locus substitution rate variation. Our implementation of this model in `MCMC_SEQ` was used to estimate the rates from sequence alignments simulated using the same model, from each of the two exemplar phylogenies (a species tree and a species network with one reticulation). Rates along the diagonal exactly match the true rates used for simulation.

### 4.3 Reanalysis of empirical data

#### 4.3.1 Test statistics support reticulate evolution within *Anopheles*

We reanalyzed previously studied *Anopheles* mosquito genomes (Fontaine et al., 2015) using both existing statistical approaches and the species triplet test approach. By conducting Pearson’s, G and Monte Carlo estimation tests on the full set of species, we found the result indicated the data did not fit the null hypothesis (*p* < 0.001). By calculating test statistics on triplets of taxa, all three multiple-testing error correction methods rejected the null hypothesis regardless of the test statistics used. This result suggests that introgression exists among the species we have selected in this Anopheles data set.

#### 4.3.2 Introgression between some *Heliconius* species is no longer supported

We analyzed subsets of an empirical data set of *Heliconius* butterfly genomes (Edelman et al., 2019) using Pearson’s, G and Monte Carlo estimation tests on the subset of *Heliconius melpomene*, *H. hecalesia* and subspecies *H. erato demophoon*. We found the data was not incompatible with the MSC (*p* = 0.81 for Pearson’s test and G test, *p* = 0.98 for Monte Carlo estimation). For the subset of species *H. timareta*, *H. melpomene* and *H.numata*, the tests did not reject the MSC (*p* = 0.99 for Pearson’s test and G test, *p* = 1 for Monte Carlo estimation). This result indicates that no introgression was detected in any of these two subsets of species in this *Heliconius* data.

We analyzed the subsets using the same phylogenetic network inference methods as for our simulation study. The log-likelihood curve of phylogenetic networks inferred using the summary method `InferNetwork_ML` for the species *H. melpomene*, *H. hecalesia* and subspecies *H. erato demophoon* was almost horizontal without a noticeable shoulder. Given the robustness of summary methods at predicting reticulation number, this result suggests that the true number of reticulations for this subset is zero (Figure 12E).

**Figure 12:**
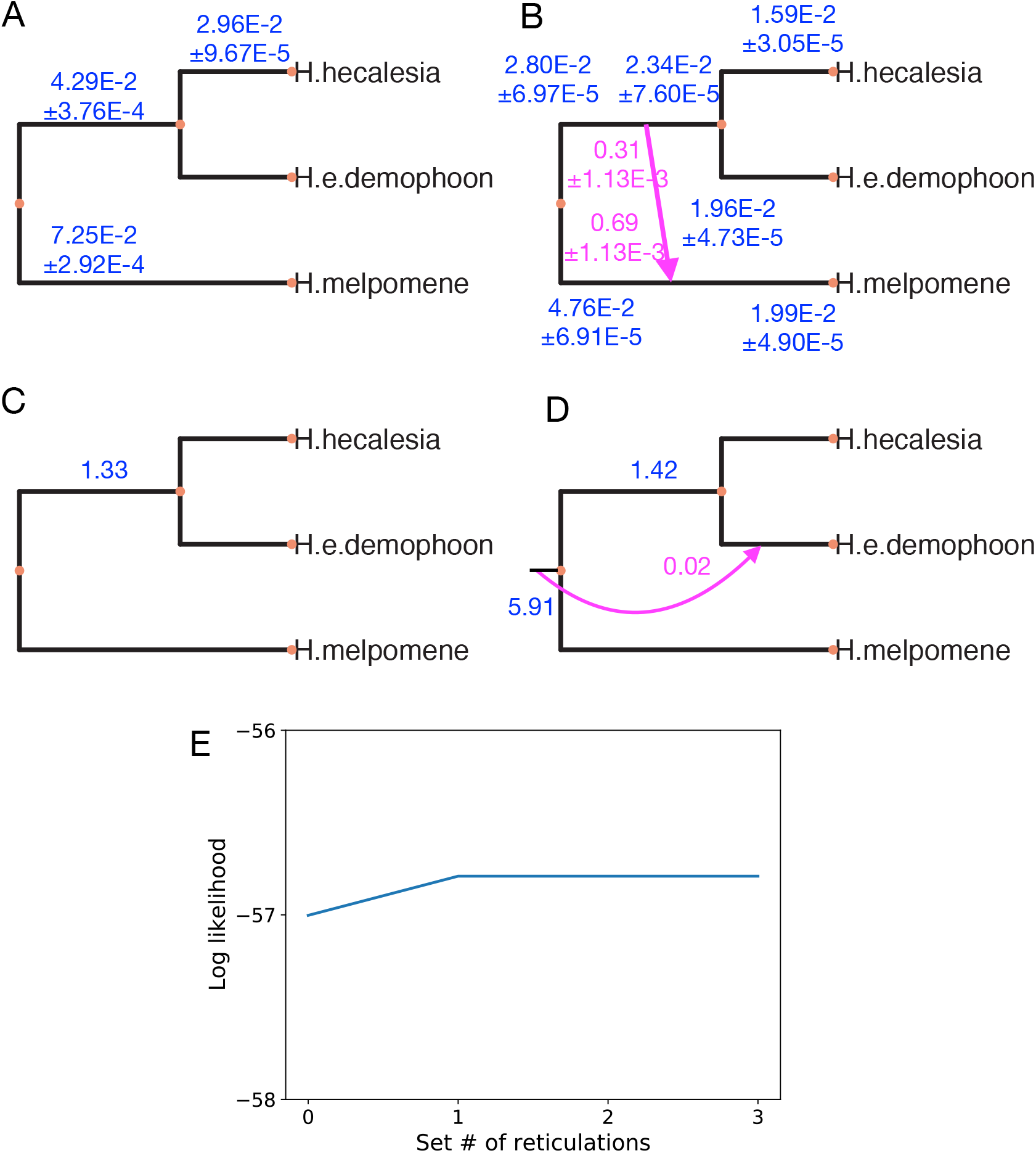
Phylogenetic network analysis of *Heliconius* data sets. Species networks were inferred using the full Bayesian method `MCMC_SEQ` (A and B) and the maximum likelihood summary method `InferNetwork_ML` (C and D), with *Heliconius* species restricted to *H. hecalesia*, *H. erato demophoon*, and *H. melpomene*. Phylogenetic inference was performed using the Dirichlet rates model of per-locus rate variation (A) or a single rate model (B). The posterior expectation and standard deviation of branch lengths are given in expected substitutions per site (blue). The posterior expectation and standard deviation of inheritance probabilities are also shown (pink). The maximum number of reticulations was set to zero (C), or up to 3 (D). Gene trees used as input were inferred using `IQ-TREE`. Estimated branch lengths are given in coalescent units (blue). Inheritance probabilities are also shown (pink). (E) Log-likelihood increase of *Heliconius* species networks identified using `InferNetworks_ML`.

When using the full Bayesian method `MCMC_SEQ` with the DEO implementation of the DR model enabled, the *maximum a posteriori* phylogeny was a tree without reticulations and an identical topology to the maximum likelihood method with the number of reticulations was set to zero (Figure 12A and C). However, when the SR model was used for inference, the full Bayesian method inferred gene flow after speciation from the ancestor of *H. hecalesia* and subspecies *H. erato demophoon* into the *H. melpomene* lineage (Figure 12B). This was different from the reticulation inferred by `InferNetwork_ML` when one reticulation was allowed, although as mentioned reticulations are unsupported by the log-likelihood curve (Figure 12D).

Substantial variation in substitution rates was inferred when using the DR model (Figure S4). Given this observation, the failure to reject statistical tests of fit to a species tree, and the trend we observed in the simulation study where using the SR model leads to the inference of spurious reticulations when rate heterogeneity is present, we suggest this apparent gene flow is an artifact of model misspecification.

The inference of spurious reticulations does not always manifest. We analyzed another subset of three species *H. timareta*, *H. melpomene* and *H. numata*, and no reticulations were inferred by the full Bayesian method regardless of whether the SR or DR models were used for inference (Figure 13).

**Figure 13:**
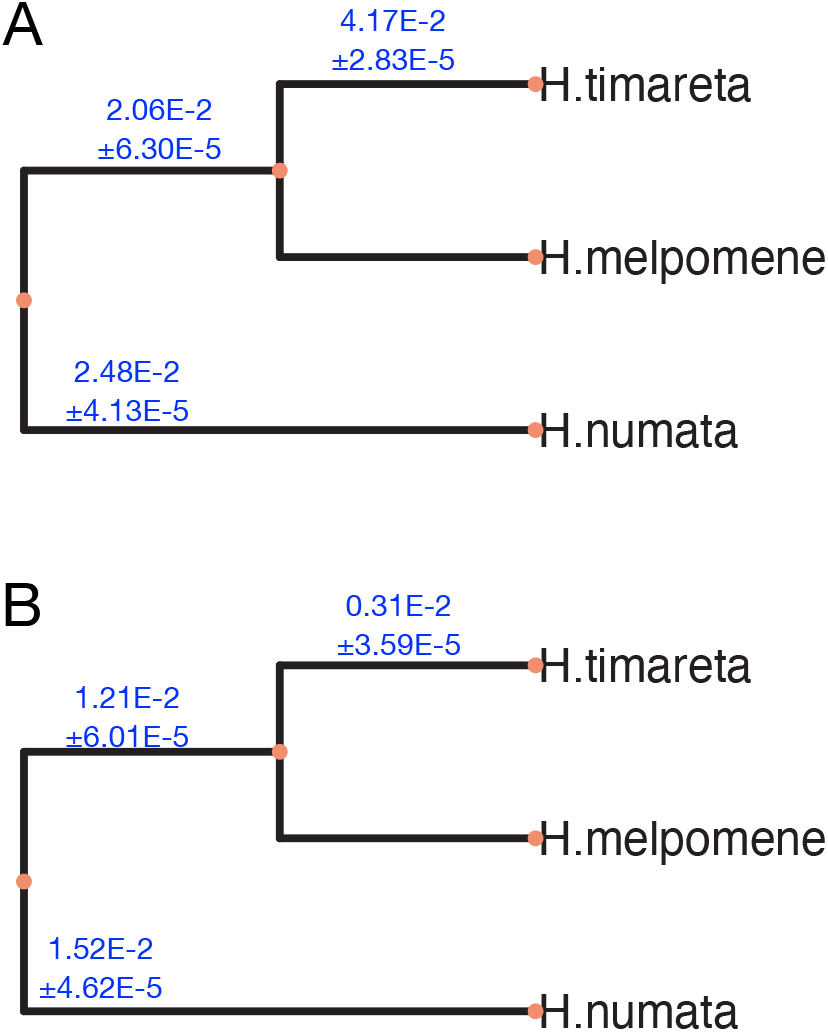
*Heliconius* species networks inferred using the full Bayesian method `MCMC_SEQ` for an alternative subset of taxa. Phylogenetic inference was performed using the Dirichlet rates model of per-locus rate variation (A) or a single rate model (B), from sequence data extracted from a whole genome alignment of *Heliconius* species, pruned to *H. timareta*, *H. melpomene* and *H. numata*. The posterior expectation and standard deviation of branch lengths are given in expected substitutions per site (blue).

In spite of variation in substitution rates also being inferred for this subset (Figure S9), this is in agreement with the log-likelihood curve and topology inferred using the summary method (Figure S7A, S8).

### 4.4 Accounting for rate variation increases convergence time

MCMC chains were run on the NOTS compute cluster at Rice University, which uses a heterogenous mix of compute nodes with four different Intel Xeon CPUs. The specific models and clock speeds are the E5-2650 v2 at 2.6GHz, the E5-2650 v4 at 2.2GHz, the Gold 6126 at 2.6GHz, and the Gold 6230 at 2.1GHz. The individual compute node used each time for a chain that was initialized or resumed was determined by the cluster scheduling software, so reported running times are reflective of a distribution of hardware rather than any particular system.

Convergence of `MCMC_SEQ` was slower when the DR model was used. When the DEO was enabled to analyze simulated data under a DR model, the unnormalized log-posterior density effective sample size (ESS) accumulation was anywhere from 63% to 17% the rate of ESS accumulation when DEO was disabled. For the subsets of *Heliconius* taxa, employing the DR model resulted in ESS accumulation being 62% or 4% the rate of ESS accumulation when employing an SR model (Table 1).

**Table 1:** Running time and ESS on simulated and empirical butterfly data for MCMC chains.

|  |  | SR inference |  |  | DR inference |  |  |
| --- | --- | --- | --- | --- | --- | --- | --- |
|  |  | ESS | Runtime<sup>a</sup> | ESS per hour | ESS | Runtime<sup>a</sup> | ESS per hour |
| Tree | SR Data | 460.19 | 24.25 | 18.98 | 289.47 | 24.18 | 11.97 |
|  | DR Data | 1169.53 | 37.05 | 31.57 | 180.14 | 25.49 | 7.07 |
| Network | SR Data | 641.46 | 25.66 | 25.00 | 269.93 | 26.31 | 10.26 |
|  | DR Data | 1107.49 | 29.35 | 37.73 | 166.41 | 26.63 | 6.25 |
| Heliconius | MelHecErd<sup>b</sup> | 11737 | 142.18 | 82.55 | 436 | 124.62 | 3.50 |
|  | TimMelNum<sup>c</sup> | 2920 | 163.63 | 17.85 | 1714 | 155.37 | 11.03 |

<sup>a</sup> Runtime was measured in hours of walltime
<sup>b</sup> The subset of *Heliconius melpomene*, *H. hecalesia* and subspecies *H. erato demophoon*
<sup>c</sup> The subset of *H. timareta*, *H. melpomene* and *H. numata*

## 5 Discussion

Model misspecification is a persistent issue in phylogenetic inference. A major area of misspecification is where gene flow occurs after speciation, which is not accounted for by simple species *tree* models. The authors of this paper, and other developers of methods for systematics, have made progress in relaxing this assumption so that inferred species phylogenies can incorporate gene flow through a species *network* model. However, we have shown that in resolving one area of misspecification, we have made the inference of species phylogenies more susceptible to misspecified substitution models.

We set out to understand the effect of model misspecification on phylogenetic network inference. Via simulation, when the model is misspecified by GTEE, we have found the test statistics are prone to have large false positives when the true model is the MSC, though these methods are good at assessing that the MSC is unfit to data generated under the MSNC. We investigated a triplet-based approach using multiple-testing error correction that showed very promising results. Although useful, this triplet-based approach we proposed has limitations, as it is restricted to using species tree as the null hypothesis, rather than a species network, due to the lack of identifiability of parameters in a three-taxon network from the frequencies of gene tree topologies. A three-taxon network, or trinet, has four internal branches, each associated with a parameter that reflects its length, plus an inheritance probability parameter associated with the reticulation edges. However, there are only three gene tree topologies, which results in a “more variables than equations” scenario, and hence the lack of trinet parameter identifiability.

Another source that misspecifies the model is the per-locus rate heterogeneity, which can cause spurious reticulations using full methods when it is not accounted for. This observation was not unexpected, as these methods interpret for any signal deviating from the MSC as evidence for introgression. Adding the DEO restores the inherent ability of full methods to identify the number of reticulations with appropriate priors. While we made the network inference more accurate by sampling more parameters to account for rate heterogeneity practically, there is a trade-off between model complexity and scalability for Bayesian methods (Fisher et al., 2022).

The accurate inference of gene trees is also harmed by this misspecification, particularly the inference of their branch lengths. For researchers interested in the patterns of substitution rate variation, we have also shown that these rates may to an extent be inferred using the DR model, even in the presence of reticulate evolution. On the other hand, when the true generative process does not incorporate rate variation, using a model for inference that allows for rate variation does not have a substantial negative impact on accuracy. Because of this asymmetry in outcomes, we recommend that species networks should be inferred using the DR or similar model whenever substitution rate variation is possible.

Through our empirical study, we have shown that this misspecification has likely caused spurious inferences when applied to real taxonomic systems. The inference of spurious reticulations may lead to incorrect conclusions concerning patterns of reticulation in evolution. This further supports our recommendation that rate variation should be accounted for whenever it may be present. To facilitate our recommendation, we have implemented the DR model in `MCMC_SEQ`, one of the most popular methods for species network inference.

Finally, determining the number of reticulations in summary phylogenetic network inference methods is expected to be more complicated when GTEE exists (Braun et al., 2019). When using maximum likelihood methods, information criteria AIC, BIC have been used to select the number of reticulations, but turn out to be ineffective (Yu et al., 2014). Similarly, the “elbow” approach was suggested to be applied to both parsimony and (pseudo-)likelihood methods by finding the number of reticulations after which the score (the number of deep coalescence events or likelihood) grows very slowly (Cao et al., 2023). But this method requires visualization and manual inspection.

## 6 Conclusions

Phylogenetic networks extend the phylogenetic tree model by allowing for nodes with two parents in order to capture the possibility of reticulations in evolutionary history. Consequently, phylogenetic networks are a richer model and in fact they can be arbitrarily complex since the number of reticulations that can be added to a phylogenetic network with a fixed number of taxa is unbounded in theory. The richness of the model can lead to erroneous inferences in practice, as any deviations from a tree-based null model, even when these deviations are not caused by reticulation, will be interpreted as reticulation by phylogenetic network inference methods. In this paper, we investigated two important sources of model misspecification, namely GTEE and substitution rate heterogeneity across loci. The former impacts methods that utilize gene tree estimates as input, whereas the latter impacts full Bayesian approaches that utilize multi-locus sequence alignments. We found that GTEE has an outsized impact on test statistics aimed at determining whether the evolutionary history is treelike or not, and showed that running the test on three-taxon subsets and combining the results could significantly ameliorate the problem. We further showed that accounting for variation in per-locus substitution rates could significantly improve the reliability of full Bayesian inference methods.

## 7 Acknowledgements

We are grateful to Zejian Liu for discussions about the statistical tests. This work was supported in part by NSF grants CCF-1514177, CCF-1800723, CNS-2131294 and DBI-2030604 to L.N., and by DMS-2153704 to H.A.O.

## References

Benjamini, Y., & Hochberg, Y. (1995). Controlling the false discovery rate: A practical and powerful approach to multiple testing. Journal of the Royal Statistical Society Series B (Statistical Methodology), 57, 289–300. (https://doi.org/10.1111/j.2517-6161.1995.tb02031.x

Bradley, D. R., Bradley, T., McGrath, S. G., & Cutcomb, S. D. (1979). Type I error rate of the chi-square test in independence in R × C tables that have small expected frequencies. Psychological Bulletin, 86, 1290–1297. https://doi.org/10.1037/0033-2909.86.6.1290

Braun, E. L., Cracraft, J., & Houde, P. (2019). Resolving the avian tree of life from top to bottom: The promise and potential boundaries of the phylogenomic era. In R. H. Kraus (Ed.), Avian genomics in ecology and evolution (pp. 151–210). Springer. https://doi.org/10.1007/978-3-030-16477-5%5C_6

Cai, R., & Ané, C. (2021). Assessing the fit of the multi-species network coalescent to multi-locus data. Bioinformatics, 37, 634–641. https://doi.org/10.1093/bioinformatics/btaa863

Cao, Z., Liu, X., Ogilvie, H. A., Yan, Z., & Nakhleh, L. (2023). Practical aspects of phylogenetic network analysis using PhyloNet [in press]. In L. S. Kubatko & L. L. Knowles (Eds.), Species tree inference. Princeton University Press. https://www.biorxiv.org/content/10.1101/746362v1

Cao, Z., Zhu, J., & Nakhleh, L. (2019). Empirical performance of tree-based inference of phylogenetic networks. 19th International Workshop on Algorithms in Bioinformatics (WABI). https://doi.org/10.4230/LIPIcs.WABI.2019.21

Davidson, R., Vachaspati, P., Mirarab, S., & Warnow, T. (2015). Phylogenomic species tree estimation in the presence of incomplete lineage sorting and horizontal gene transfer. BMC Genomics, 16, S1. https://doi.org/10.1186/1471-2164-16-S10-S1

Degnan, J. H., & Rosenberg, N. A. (2009). Gene tree discordance, phylogenetic inference and the multispecies coalescent. Trends in Ecology & Evolution, 24, 332–340. https://doi.org/10.1016/j.tree.2009.01.009

Degnan, J. H., & Salter, L. A. (2005). Gene tree distributions under the coalescent process. Evolution, 59, 24–37. https://doi.org/10.1111/j.0014-3820.2005.tb00891.x

Drummond, A. J., & Rambaut, A. (2007). BEAST: Bayesian evolutionary analysis by sampling trees. BMC Evolutionary Biology, 7, 214. https://doi.org/10.1186/1471-2148-7-214

Duret, L., & Mouchiroud, D. (2000). Determinants of substitution rates in mammalian genes: Expression pattern affects selection intensity but not mutation rate. Molecular Biology and Evolution, 17, 68–70. https://doi.org/10.1093/oxfordjournals.molbev.a026239

Edelman, N. B., Frandsen, P. B., Miyagi, M., Clavijo, B., Davey, J., Dikow, R. B., García-Accinelli, G., Van Belleghem, S. M., Patterson, N., Neafsey, D. E., et al. (2019). Genomic architecture and introgression shape a butterfly radiation. Science, 366, 594–599. https://doi.org/10.1126/science.aaw2090

Fisher, A. A., Hassler, G. W., Ji, X., Baele, G., Suchard, M. A., & Lemey, P. (2022). Scalable Bayesian phylogenetics. Philosophical Transactions of the Royal Society B: Biological Sciences, 377, 20210242. https://doi.org/10.1098/rstb.2021.0242

Flouri, T., Jiao, X., Rannala, B., & Yang, Z. (2020). A Bayesian implementation of the multispecies coalescent model with introgression for phylogenomic analysis. Molecular Biology and Evolution, 37, 1211–1223. https://doi.org/10.1093/molbev/msz296

Fontaine, M. C., Pease, J. B., Steele, A., Waterhouse, R. M., Neafsey, D. E., Sharakhov, I. V., Jiang, X., Hall, A. B., Catteruccia, F., Kakani, E., Mitchell, S. N., Wu, Y.-C., Smith, H. A., Love, R. R., Lawniczak, M. K., Slotman, M. A., Emrich, S. J., Hahn, M. W., & Besansky, N. J. (2015). Extensive introgression in a malaria vector species complex revealed by phylogenomics. Science, 347, 1258524. https://doi.org/10.1126/science.1258524

Green, R. E., Krause, J., Briggs, A. W., Maricic, T., Stenzel, U., Kircher, M., Patterson, N., Li, H., Zhai, W., Fritz, M. H.-Y., Hansen, N. F., Durand, E. Y., Malaspinas, A.-S., Jensen, J. D., Marques-Bonet, T., Alkan, C., Prafer, K., Meyer, M., Burbano, H. A.,... Paabo, S. (2010). A draft sequence of the Neandertal genome. Science, 328, 710–722. https://doi.org/10.1126/science.1188021

Hahn, M. W., & Hibbins, M. S. (2019). A three-sample test for introgression. Molecular Biology and Evolution, 36, 2878–2882. https://doi.org/10.1093/molbev/msz178

Härdle, W., Horowitz, J., & Kreiss, J.-P. (2003). Bootstrap methods for time series. International Statistical Review, 71, 435–459. https://www.jstor.org/stable/1403897

Heled, J., & Drummond, A. J. (2009). Bayesian inference of species trees from multilocus data. Molecular Biology and Evolution, 27, 570–580. https://doi.org/10.1093/molbev/msp274

Hochberg, Y. (1988). A sharper Bonferroni procedure for multiple tests of significance. Biometrika, 75, 800–802. https://doi.org/10.1093/biomet/75.4.800

Hosmane, B. (1987). Improved likelihood ratio test for multinomial goodness of fit. Communications in Statistics—Theory and Methods, 16, 3185–3198. https://doi.org/10.1080/03610928708829566

Hudson, R. R. (2002). Generating samples under a Wright-Fisher neutral model of genetic variation. Bioinformatics, 18, 337–338. https://doi.org/10.1093/bioinformatics/18.2.337

Jukes, T. H., & Cantor, C. R. (1969). Evolution of protein molecules. In H. N. Munro (Ed.), Mammalian protein metabolism (pp. 21–132). Academic Press. https://doi.org/10.1016/B978-1-4832-3211-9.50009-7

Larntz, K. (1978). Small-sample comparisons of exact levels for chi-squared goodness-of-fit statistics. Journal of the American Statistical Association, 73, 253–263. https://doi.org/10.1080/01621459.1978.10481567

Marcussen, T., Sandve, S. R., Heier, L., Spannagl, M., Pfeifer, M., The International Wheat Genome Sequencing Consortium, Jakobsen, K. S., Wulff, B. B., Steuernagel, B., Mayer, K. F., & Olsen, O.-A. (2014). Ancient hybridizations among the ancestral genomes of bread wheat. Science, 345, 1250092. https://doi.org/10.1126/science.1250092

Morales-Briones, D. F., Kadereit, G., Tefarikis, D. T., Moore, M. J., Smith, S. A., Brockington, S. F., Timoneda, A., Yim, W. C., Cushman, J. C., & Yang, Y. (2021). Disentangling sources of gene tree discordance in phylogenomic data sets: Testing ancient hybridizations in Amaranthaceae s.l. Systematic Biology, 70, 219–235. https://doi.org/10.1093/sysbio/syaa066

Nakhleh, L. (2009). A metric on the space of reduced phylogenetic networks. IEEE/ACM Transactions on Computational Biology and Bioinformatics, 7, 218–222. https://doi.org/10.1109/TCBB.2009.2

Nakhleh, L. (2013). Computational approaches to species phylogeny inference and gene tree reconciliation. Trends in Ecology & Evolution, 28, 719–728. https://doi.org/10.1016/j.tree.2013.09.004

Nguyen, L.-T., Schmidt, H. A., Von Haeseler, A., & Minh, B. Q. (2015). IQ-TREE: A fast and effective stochastic algorithm for estimating maximum-likelihood phylogenies. Molecular Biology and Evolution, 32, 268–274. https://doi.org/10.1093/molbev/msu300

Pearson, K. (1900). On the criterion that a given system of deviations from the probable in the case of a correlated system of variables is such that it can be reasonably supposed to have arisen from random sampling. The London, Edinburgh, and Dublin Philosophical Magazine and Journal of Science, 50, 157–175. https://doi.org/10.1080/14786440009463897

Rambaut, A., & Grass, N. C. (1997). Seq-Gen: An application for the Monte Carlo simulation of DNA sequence evolution along phylogenetic trees. Bioinformatics, 13, 235–238. https://doi.org/10.1093/bioinformatics/13.3.235

Rasmussen, M. D., & Kellis, M. (2012). Unified modeling of gene duplication, loss, and coalescence using a locus tree. Genome Research, 22, 755–765. https://doi.org/10.1101/gr.123901.111

Resin, J. (2022). A simple algorithm for exact multinomial tests [published online before print]. Journal of Computational and Graphical Statistics. https://doi.org/10.1080/10618600.2022.2102026

Robinson, D. F., & Foulds, L. R. (1981). Comparison of phylogenetic trees. Mathematical Biosciences, 53, 131–147. https://doi.org/10.1016/0025-5564(81)90043-2

Roch, S., & Snir, S. (2013). Recovering the treelike trend of evolution despite extensive lateral genetic transfer: A probabilistic analysis. Journal of Computational Biology, 20, 93–112. https://doi.org/10.1089/cmb.2012.0234

Seabold, S., & Perktold, J. (2010). Statsmodels: Econometric and statistical modeling with Python. In S. van der Walt & J. Millman (Eds.), Proceedings of the 9th Python in science conference (pp. 92–96). SciPy. https://doi.org/10.25080/Majora-92bf1922-011

Seo, T.-K. (2008). Calculating bootstrap probabilities of phylogeny using multilocus sequence data. Molecular Biology and Evolution, 25, 960–971. https://doi.org/10.1093/molbev/msn043

Simes, R. J. (1986). An improved Bonferroni procedure for multiple tests of significance. Biometrika, 73 (3), 751–754. https://doi.org/10.2307/2336545

Solís-Lemus, C., Yang, M., & Ané, C. (2016). Inconsistency of species tree methods under gene flow. Systematic Biology, 65, 843–851. https://doi.org/10.1093/sysbio/syw030

Stenz, N. W., Larget, B., Baum, D. A., & Ané, C. (2015). Exploring tree-like and non-tree-like patterns using genome sequences: An example using the inbreeding plant species Arabidopsis thaliana (l.) heynh. Systematic Biology, 64, 809–823. https://doi.org/10.1093/sysbio/syv039

Szöllősi, G. J., Tannier, E., Daubin, V., & Boussau, B. (2014). The inference of gene trees with species trees. Systematic Biology, 64, e42–e62. https://doi.org/10.1093/sysbio/syu048

Tavaré, S. (1986). Some probabilistic and statistical problems in the analysis of DNA sequences. Some Mathematical Questions in Biology—DNA Sequence Analysis, 17.

Than, C., Ruths, D., & Nakhleh, L. (2008). PhyloNet: A software package for analyzing and reconstructing reticulate evolutionary relationships. BMC Bioinformatics, 9, 322. https://doi.org/10.1186/1471-2105-9-322

Wen, D., & Nakhleh, L. (2018). Co-estimating reticulate phylogenies and gene trees from multi-locus sequence data. Systematic Biology, 67, 439–457. https://doi.org/10.1093/sysbio/syx085

Wen, D., Yu, Y., Hahn, M. W., & Nakhleh, L. (2016). Reticulate evolutionary history and extensive introgression in mosquito species revealed by phylogenetic network analysis. Molecular Ecology, 25, 2361–2372. https://doi.org/10.1111/mec.13544

Wen, D., Yu, Y., & Nakhleh, L. (2016). Bayesian inference of reticulate phylogenies under the multispecies network coalescent. PLOS Genetics, 12, e1006006. https://doi.org/10.1371/journal.pgen.1006006

Wen, D., Yu, Y., Zhu, J., & Nakhleh, L. (2018). Inferring phylogenetic networks using PhyloNet. Systematic Biology, 67, 735–740. https://doi.org/10.1093/sysbio/syy015

Woolf, B. (1957). The log likelihood ratio test (the G-test). Annals of Human Genetics, 21, 397–409. https://doi.org/10.1111/j.1469-1809.1972.tb00293.x

Yu, Y., Barnett, R. M., & Nakhleh, L. (2013). Parsimonious inference of hybridization in the presence of incomplete lineage sorting. Systematic Biology, 62, 738–751. https://doi.org/10.1093/sysbio/syt037

Yu, Y., Dong, J., Liu, K. J., & Nakhleh, L. (2014). Maximum likelihood inference of reticulate evolutionary histories. Proceedings of the National Academy of Sciences, 111, 16448–6453. https://doi.org/10.1073/pnas.1407950111

Zhang, C., Ogilvie, H. A., Drummond, A. J., & Stadler, T. (2018). Bayesian inference of species networks from multilocus sequence data. Molecular Biology and Evolution, 35, 504–517. https://doi.org/10.1093/molbev/msx307

Zhu, J., & Nakhleh, L. (2018). Inference of species phylogenies from bi-allelic markers using pseudo-likelihood. Bioinformatics, 34, i376–i385. https://doi.org/10.1093/bioinformatics/bty295

Zhu, J., Wen, D., Yu, Y., Meudt, H. M., & Nakhleh, L. (2018). Bayesian inference of phylogenetic networks from bi-allelic genetic markers. PLOS Computational Biology, 14, e1005932. https://doi.org/10.1371/journal.pcbi.1005932

Zhu, J., Yu, Y., & Nakhleh, L. (2016). In the light of deep coalescence: Revisiting trees within networks. BMC Bioinformatics, 17, 415. https://doi.org/10.1186/s12859-016-1269-1

