## Supplementary Figures for "The Impact of Model Misspecification on Phylogenetic Network Inference"

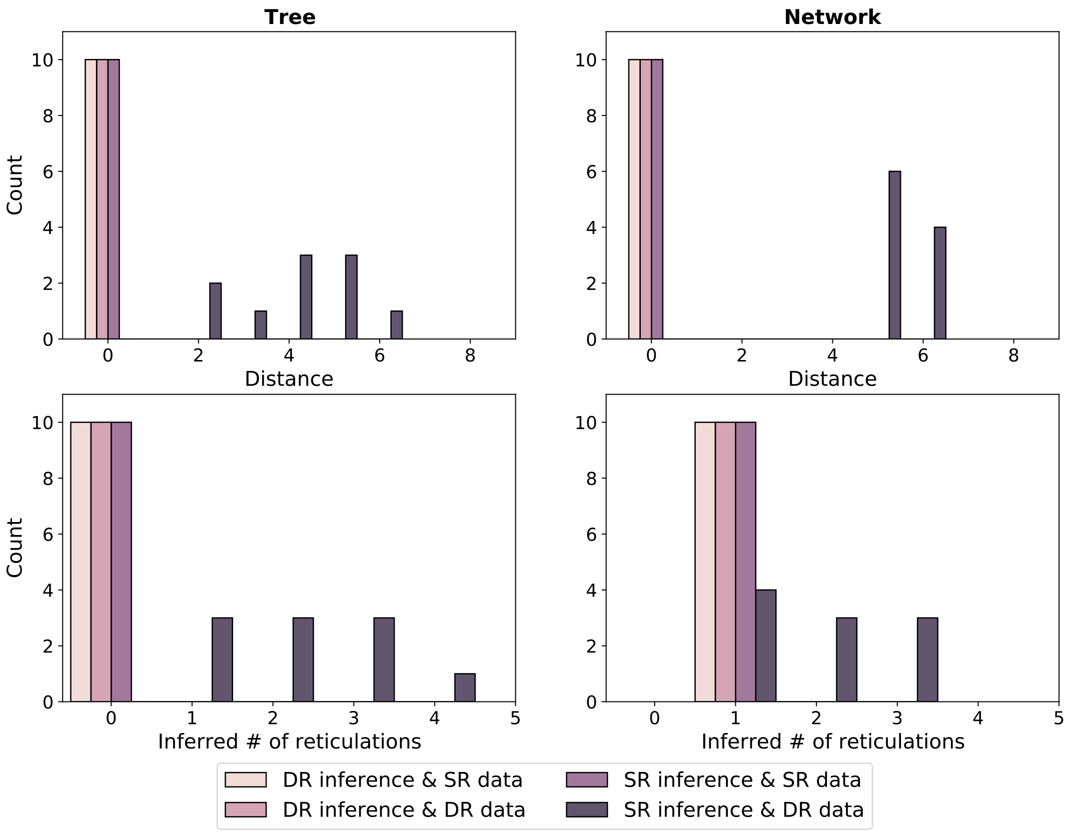


Figure S1: The effect of per-locus substitution rate variation on the accuracy of species phylogenies inferred using a full Bayesian approach. `MCMC_SEQ` was used to infer the species phylogeny together with the gene trees from multiple sequence alignments in a full Bayesian approach, under either single rate (SR) or Dirichlet rates (DR) models of per-locus substitution rate variation, and the network of the largest marginal probability was selected. Sequence alignments were also simulated under either SR or DR models. Each bar represents the number of replicates a given network distance from the true species network, for the two exemplar species phylogenies (Tree and Network).

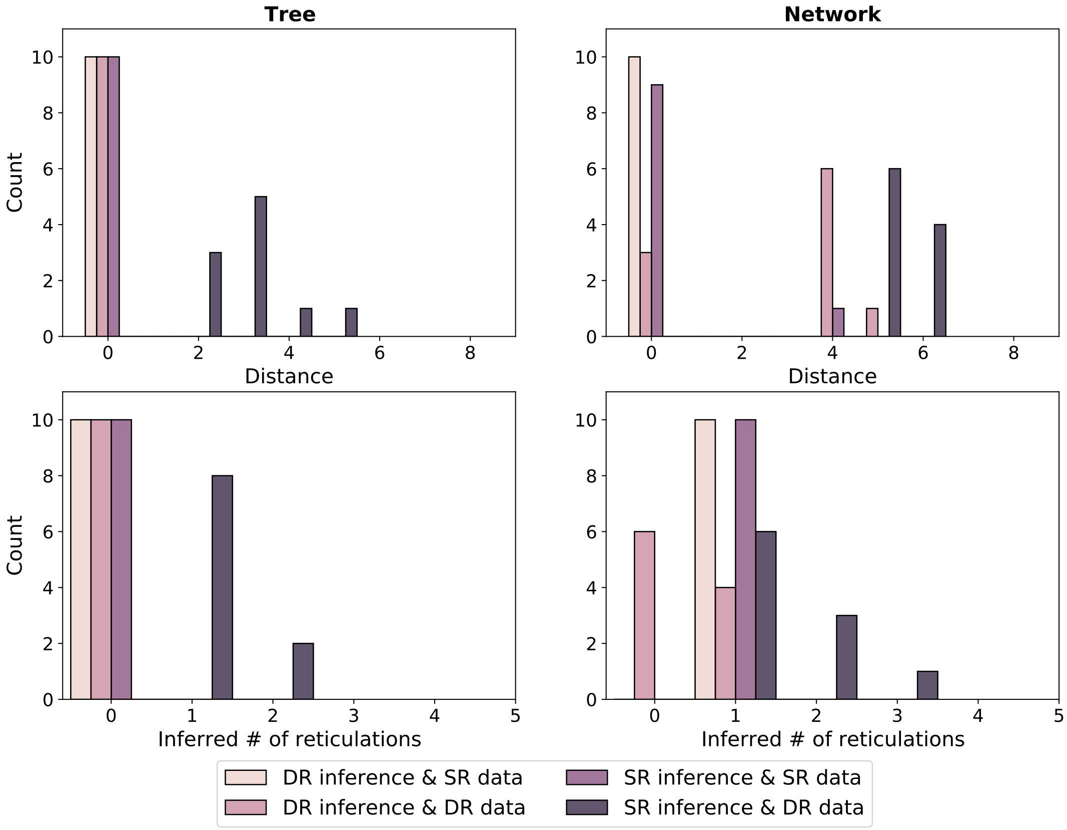


Figure S2: The effect of per-locus substitution rate variation on the accuracy of species phylogenies inferred using a full Bayesian approach. MCMC_SEQ was used to infer the species phylogeny together with the gene trees from multiple sequence alignments in a full Bayesian approach, under either single rate (SR) or Dirichlet rates (DR) models of per-locus substitution rate variation. Sequence alignments were also simulated under either SR or DR models. Result network was summarized from MAP. Each bar represents the number of replicates a given network distance from the true species network, for the two exemplar species phylogenies (Tree and Network) with with moderate ILS level and 2000nt.

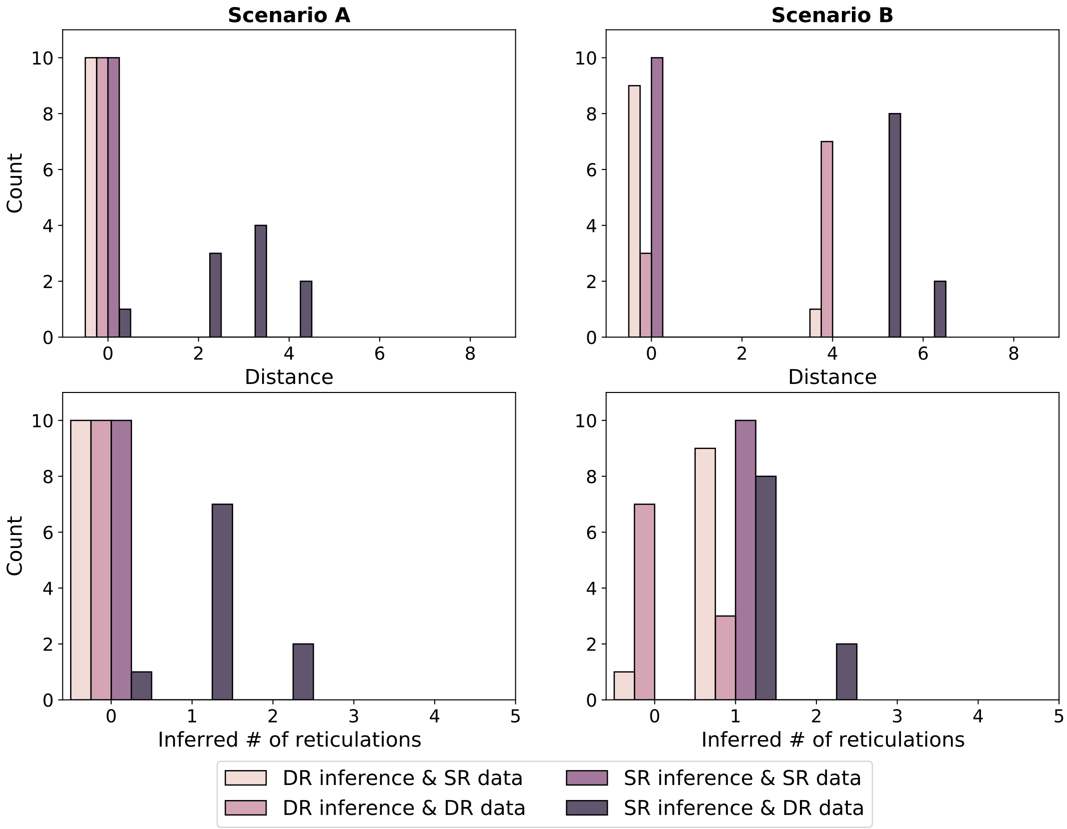


Figure S3: The effect of per-locus substitution rate variation on the accuracy of species phylogenies inferred using a full Bayesian approach. MCMC_SEQ was used to infer the species phylogeny together with the gene trees from multiple sequence alignments in a full Bayesian approach, under either single rate (SR) or Dirichlet rates (DR) models of per-locus substitution rate variation. Sequence alignments were also simulated under either SR or DR models. Result network was summarized from with the highest probability marginalized over continuous parameters. Each bar represents the number of replicates a given network distance from the true species network, for the two exemplar species phylogenies (Tree and network) with moderate ILS level and 2000nt.

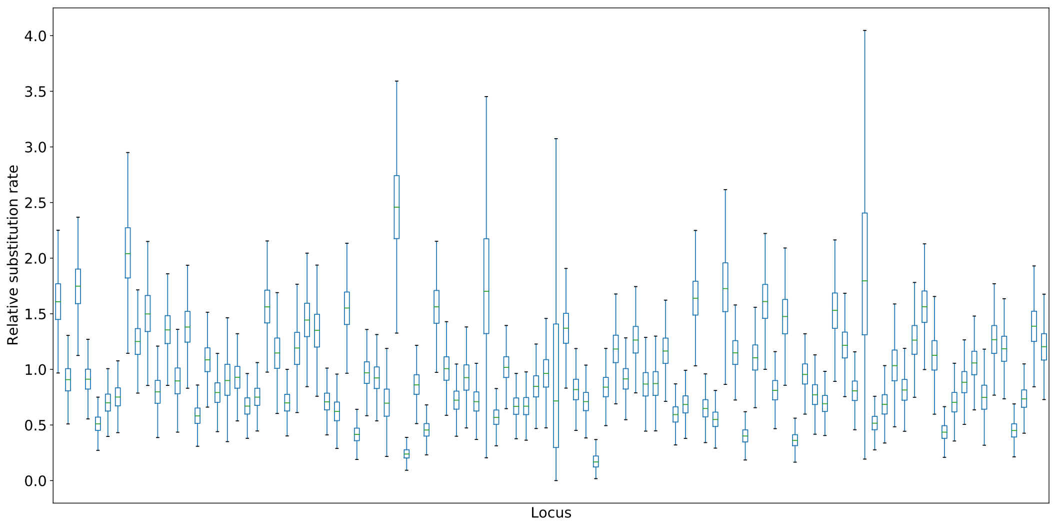


Figure S4: The inferred relative substitution rates of the butterfly dataset by `MCMC_SEQ` using DEO implementation of the DR model for species *Heliconius melpomene*, *H. erato demophoon* and *H. hecalesia*. Each box plot shows the posterior distribution of substitution rates for a separate individual locus.

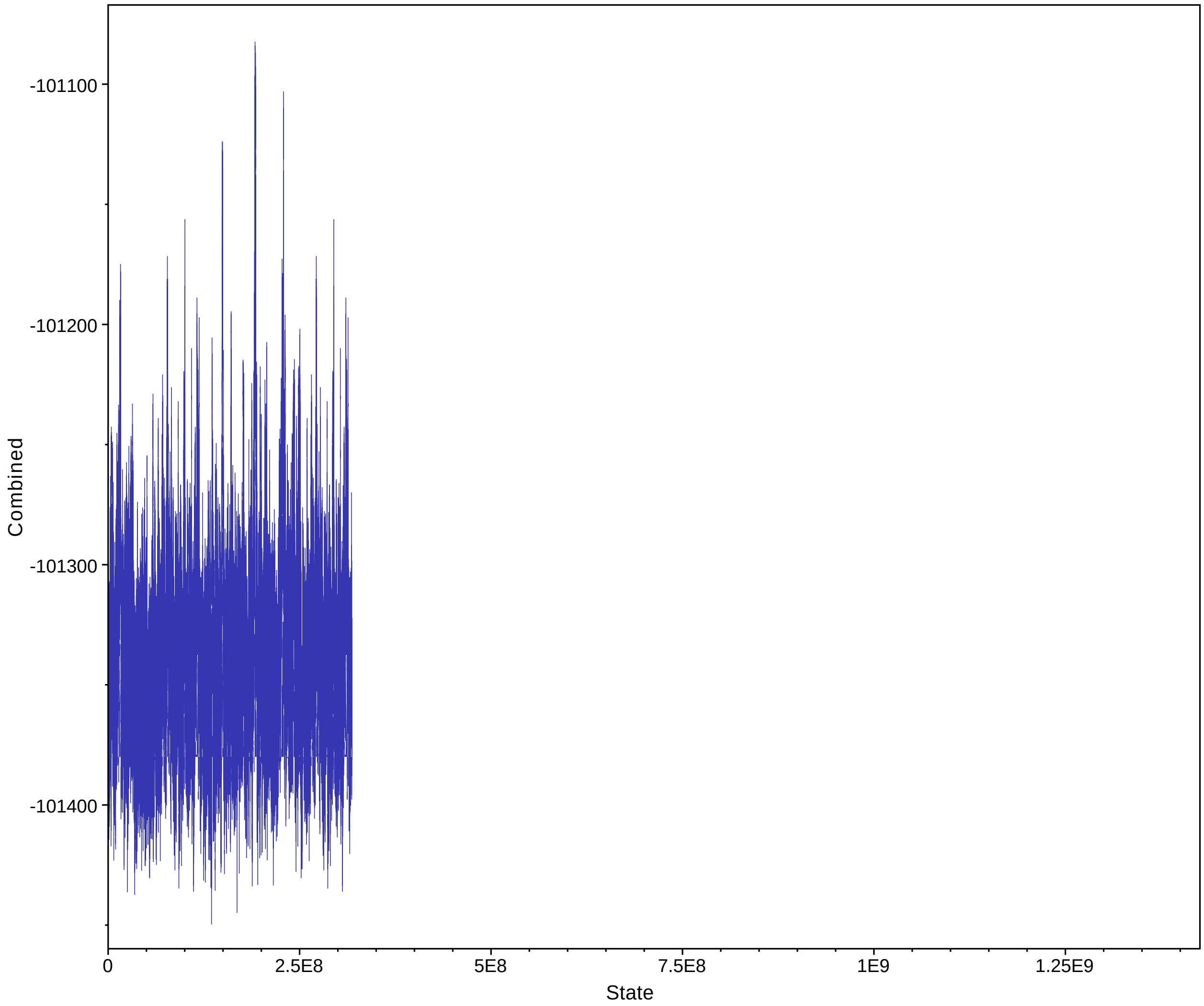


Figure S5: The posterior probability trace when running MCMC_SEQ using the DEO implementation of the DR model for species *Heliconius melpomene*, *H. erato demophoon* and *H. hecalesia*. The ESS is 436.

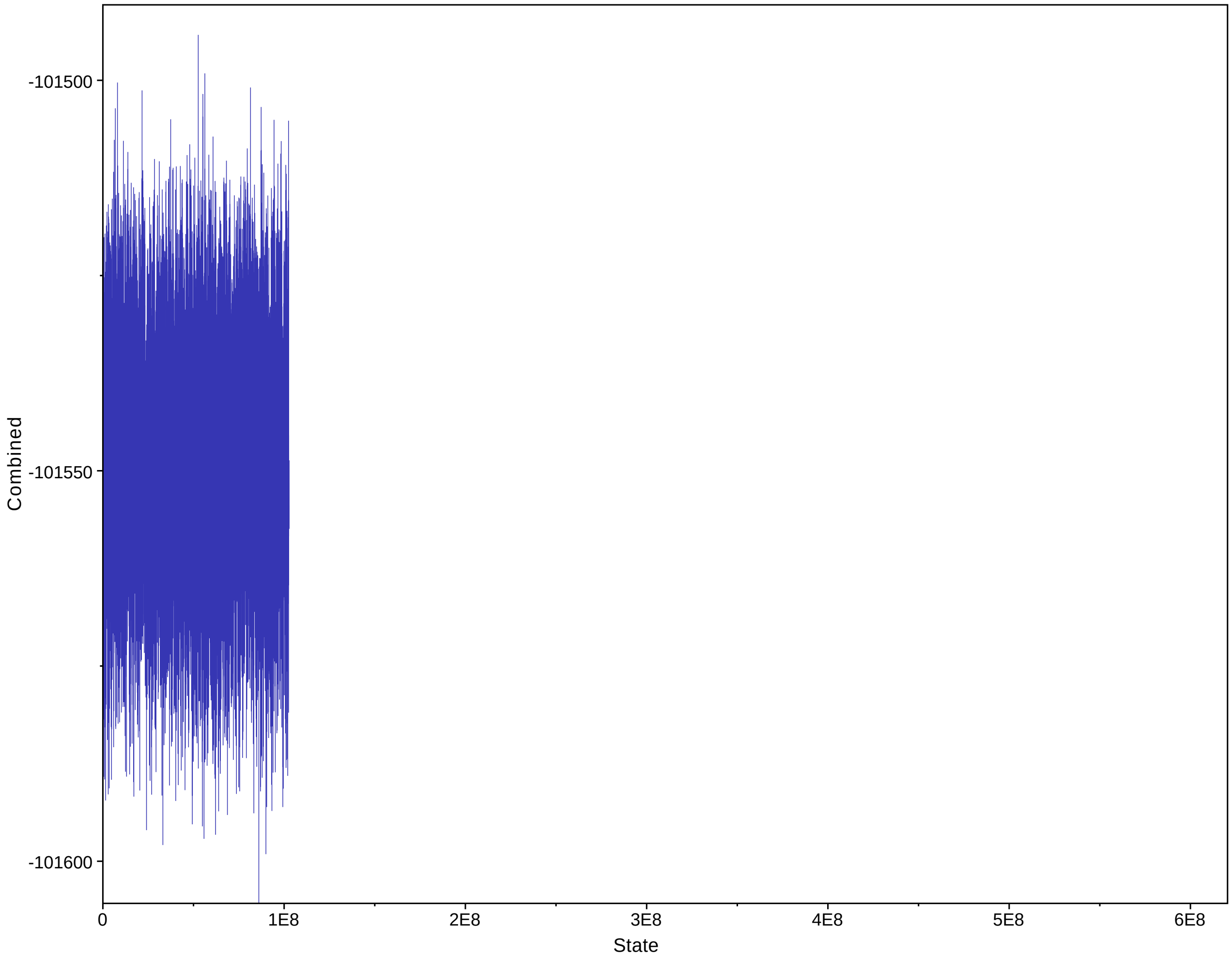


Figure S6: The posterior probability trace when running MCMC_SEQ using the SR model for species *Heliconius melpomene*, *H. erato demophoon* and *H. hecalesia*. The ESS is 11737.

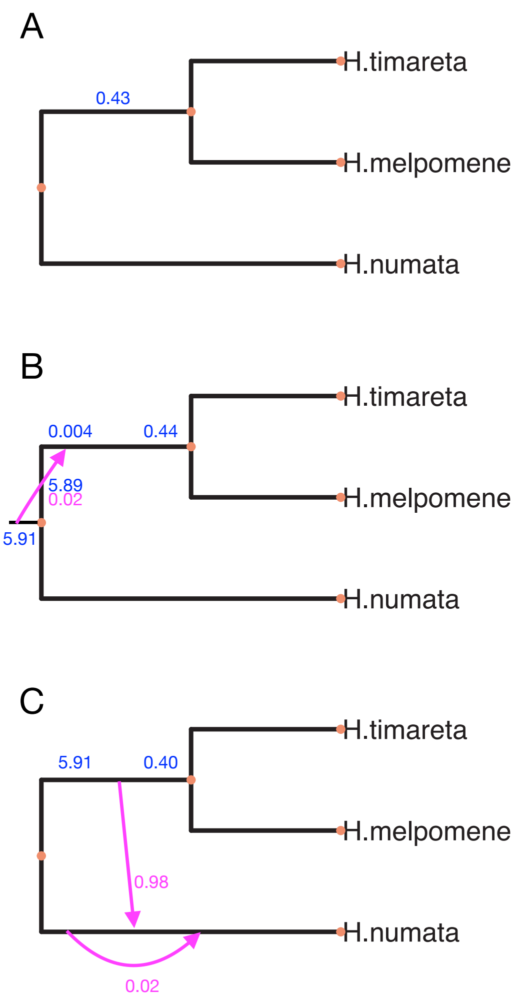


Figure S7: The species network topologies inferred by the maximum likelihood summary method `InferNetworks_ML` for species *Heliconius timareta*, *H. melpomene* and *H. numata*. A is the inferred tree when the maximum number of reticulations was set to 0 or 1. B is the inferred network when the maximum number of reticulations was set to 2. C is the inferred network when the maximum number of reticulations was set to 3.

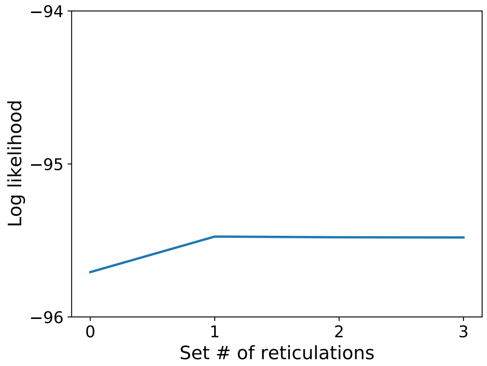


Figure S8: Log-likelihood increase of *Heliconius* species networks identified using the maximum likelihood summary method `InferNetworks_ML`. Gene trees used as input were inferred using `IQ-TREE` from sequence data extracted from an whole genome alignment of *Heliconius* species, pruned to *Heliconius timareta*, *H. melpomene* and *H. numata*.

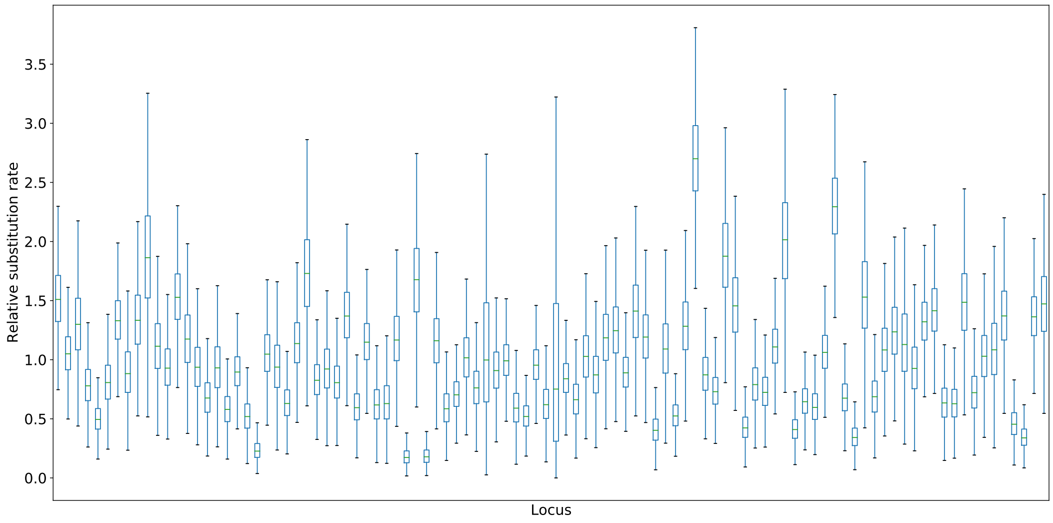


Figure S9: The inferred relative substitution rates of the butterfly dataset by `MCMC_SEQ` using the DEO implementation of the DR model for species *Heliconius timareta*, *H. melpomene* and *H. numata*.

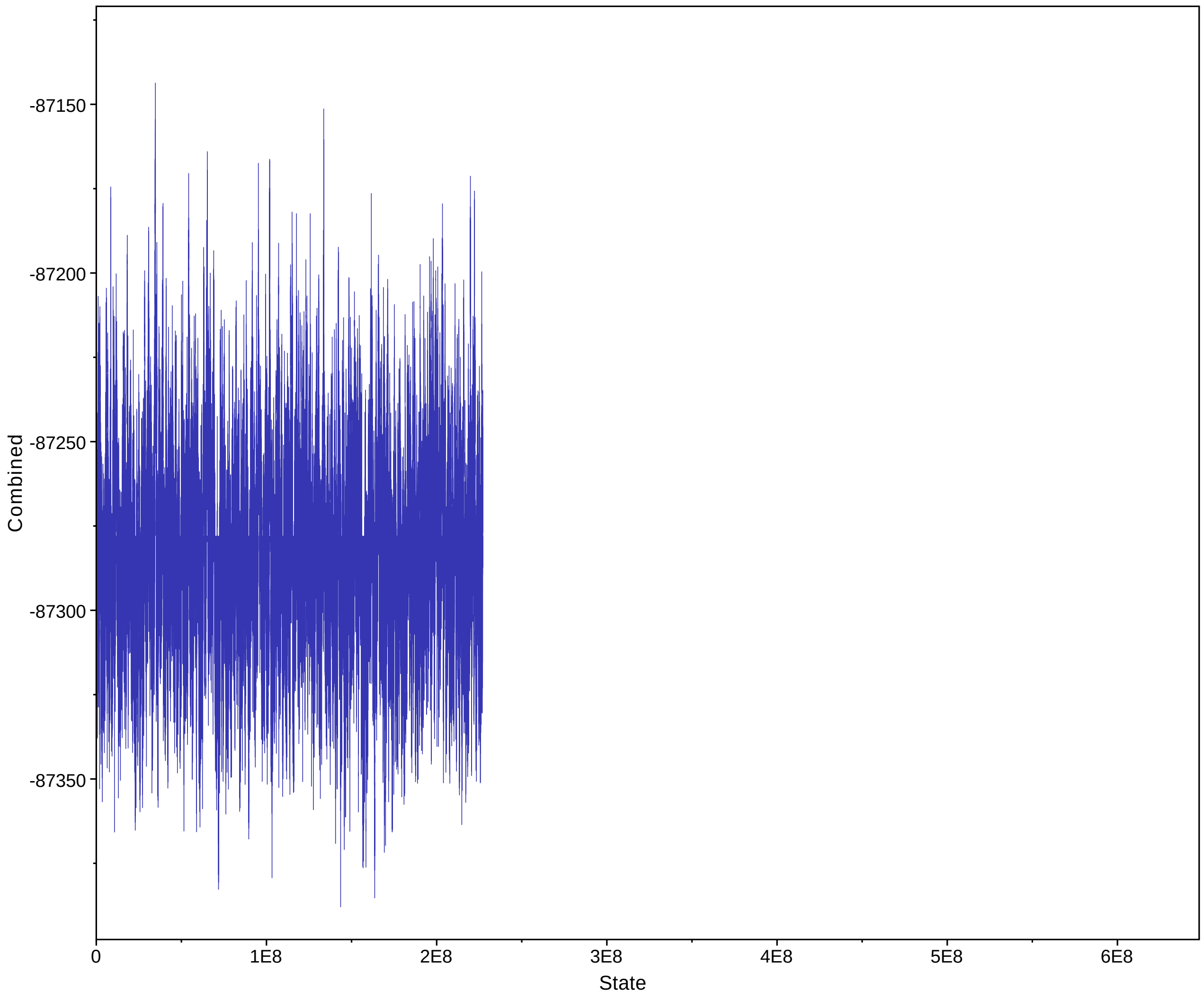


Figure S10: The posterior probability trace when running `MCMC_SEQ` using the DEO implementation of the DR model for species *Heliconius timareta*, *H. melpomene* and *H. numata*. The ESS is 1714.

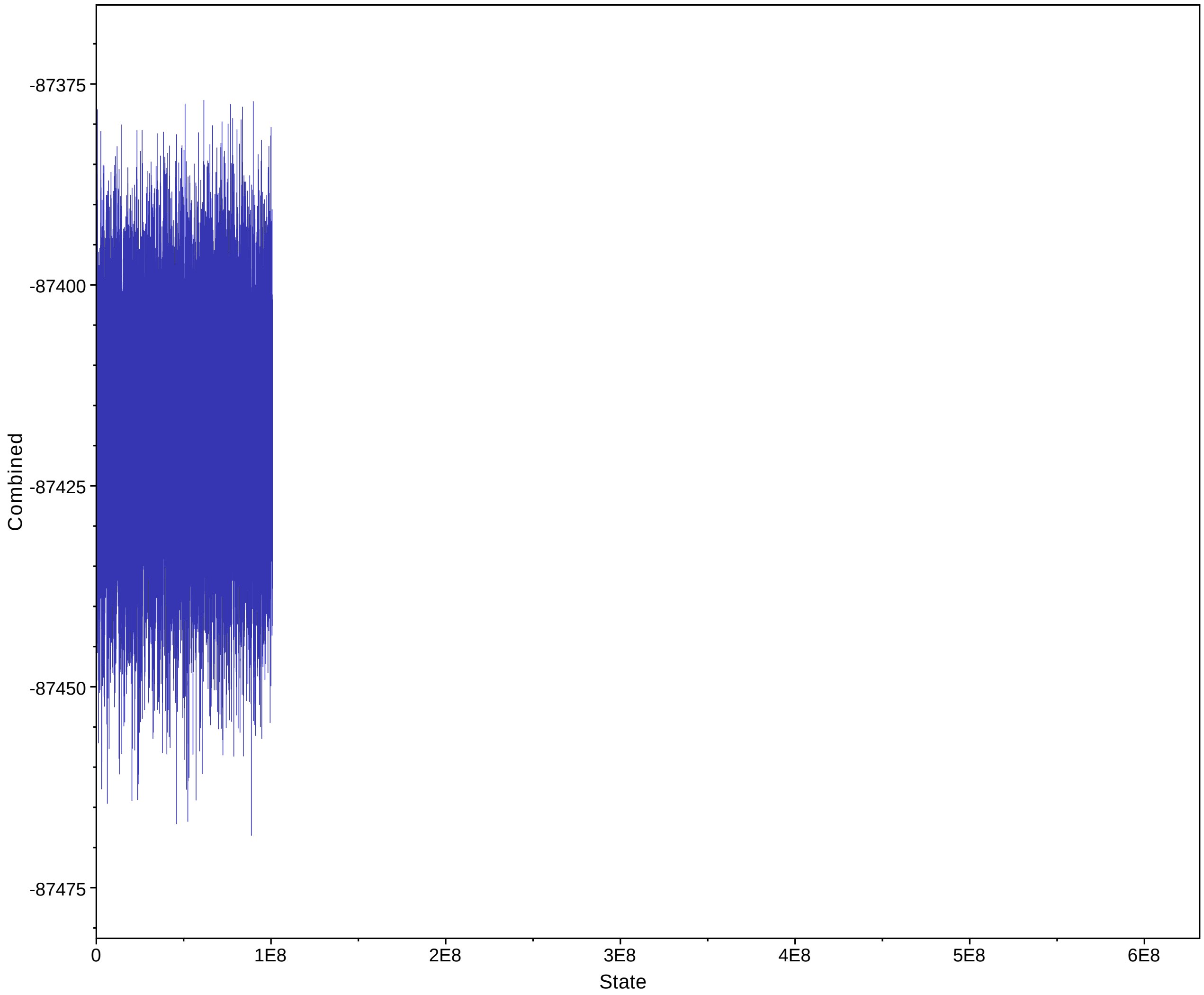


Figure S11: The posterior probability trace when running `MCMC_SEQ` using SR model for inference for species *Heliconius timareta*, *H. melpomene* and *H. numata*. The ESS is 2920.

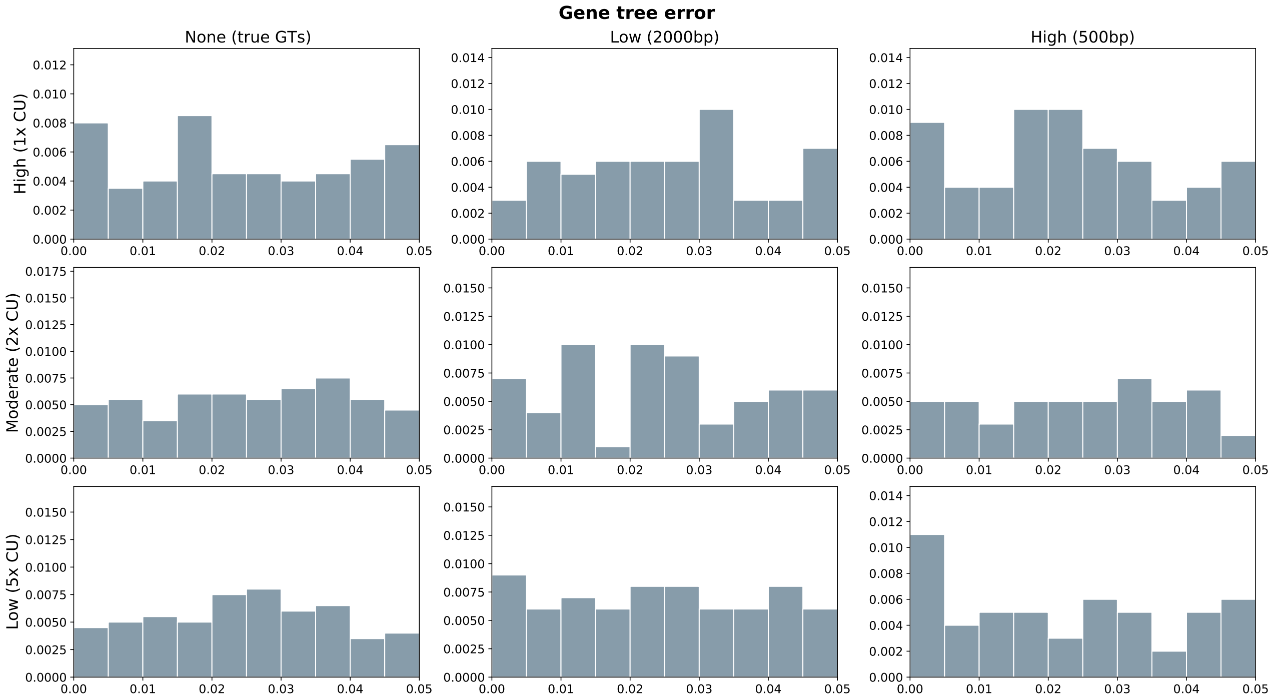


Figure S12: The histogram of triplet Pearson's $\chi^2$ test p-values that are significant for data generated under MSC.

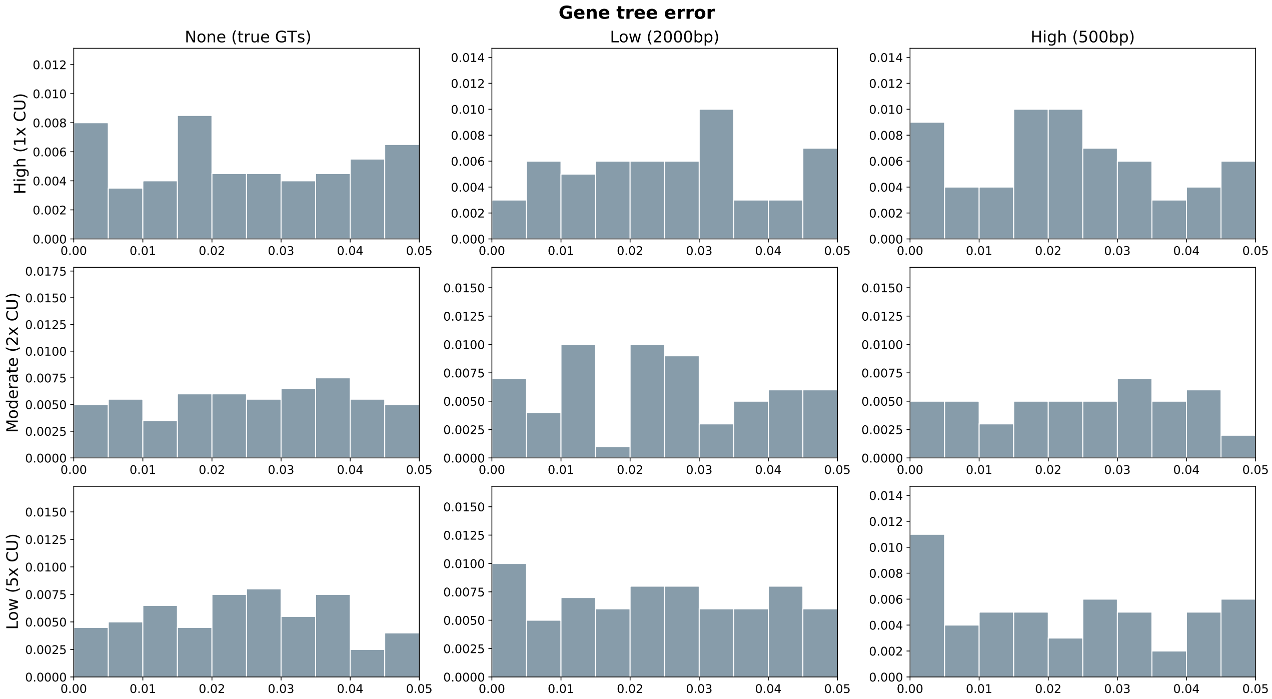


Figure S13: The histogram of triplet G-test p-values that are significant for data generated under MSC .
